# A micro-RNA is the effector gene of a classic evolutionary hotspot locus

**DOI:** 10.1101/2024.02.09.579741

**Authors:** Shen Tian, Yoshimasa Asano, Tirtha Das Banerjee, Jocelyn Liang Qi Wee, Abigail Lamb, Yehan Wang, Suriya Narayanan Murugesan, Kumiko Ui-Tei, Patricia J. Wittkopp, Antónia Monteiro

## Abstract

In Lepidoptera (butterflies and moths), the genomic region around the gene *cortex* is a ‘hotspot’ locus, repeatedly used to generate intraspecific melanic wing color polymorphisms across 100-million-years of evolution. However, the identity of the effector gene regulating melanic wing color within this locus remains unknown. Here, we show that none of the four candidate protein-coding genes within this locus, including *cortex*, serve as major effectors. Instead, a micro-RNA (miRNA), *mir-193*, serves as the major effector across three deeply diverged lineages of butterflies, and its function is conserved in *Drosophila*. In Lepidoptera, *mir-193* is derived from a gigantic long non-coding RNA, *ivory*, and it functions by directly repressing multiple pigmentation genes. We show that a miRNA can drive repeated instances of adaptive evolution in animals.

---

Hotspot loci are genomic regions that are repeatedly mutated to produce similar phenotypic variation in unrelated lineages (*1*). In Lepidoptera (butterflies and moths), a handful of evolutionarily conserved hotspot loci have been found to control striking intraspecific polymorphisms in wing color patterns (*2–13*). One of the most intensively studied loci is a genomic region containing the protein-coding gene *cortex*, associated with melanic (black and white; dark and bright) wing color pattern variations. This locus has been repeatedly used to generate adaptive wing pattern polymorphisms across a 100-million-year evolutionary history, including the textbook example of industrial melanism of the peppered moth *Biston betularia*, the iconic mimetic radiation of *Heliconius* butterflies, and the leaf wing polymorphism of oakleaf butterflies *Kallima inachus*, among others (*2, 3, 5, 7, 14*).

In previous studies, the protein-coding gene *cortex* was considered the major effector of this locus. However, in most cases, the spatial expression of *cortex* did not pre-pattern black/dark colors on the developing wings (*2, 15, 16*). Functional studies using CRISPR-Cas9 suggested that *cortex* might promote wing melanization. These studies, however, suffered from low success rates (*2, 15, 17*), hinting that hidden genomic features next to *cortex*, rather than *cortex* itself, could be causative. While previous investigations exclusively focused on protein-coding genes, non-coding RNAs were ignored, despite the presence of two micro-RNAs (miRNAs) at this locus (*18*). MiRNAs are small (20-22nt) non-coding RNAs that are well-known post-transcriptional gene regulators, but largely understudied in the genetics of morphological diversification (Fig. 1A) (*19*).

**Fig.1.**
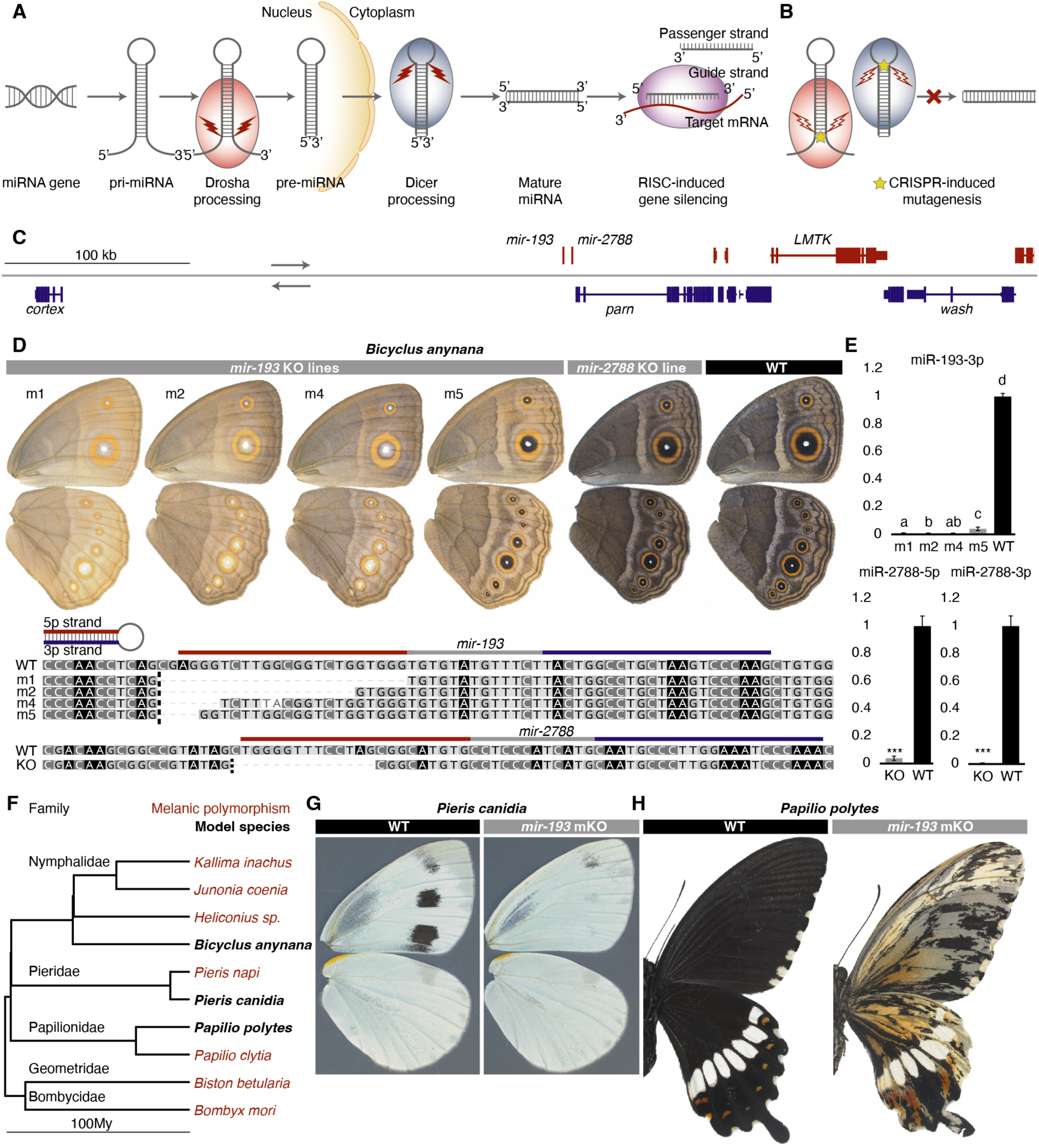
mir-193 is the major melanic color regulator in the *cortex* locus. **(A)** Biogenesis and silencing machinery of miRNAs. **(B)** Disrupting Drosha or Dicer processing sites using CRISPR-Cas9 inhibits the biogenesis of mature miRNAs. **(C)** The core *cortex* locus in *B. anynana* showing two highly conserved miRNAs and four candidate protein-coding genes. **(D)** Homozygous mutant lines of *mir-193* and *mir-2788* in *B. anynana* and their corresponding genotypes. **(E)** Expression levels of the guide strand of *mir-193*, miR-193-3p, and the two mature strands of *mir-2788*, across the corresponding mutant lines and WT. n=3-4; ns: not significant; *: p<0.05; **: p<0.01; ***: p<0.001; Expression levels with the same letter are not significantly different from each other; Error bar: SEM. **(F)** Phylogenetic placement of the three model butterfly species in the tree of lepidopterans previously mapped to the *cortex* locus. Phylogeny is from (*28*). Mosaic knock-outs (mKOs) of *mir-193* in **(G)** *Pieris canidia* and **(H)** *Papilio polytes*. Images were horizontally flipped when necessary.

## A non-coding RNA *mir-193*, but not *cortex*, is the effector of the *cortex* locus

Here, we aimed to test if miRNAs present in the *cortex* locus are the actual effectors of this locus. We began by screening the entire *cortex* locus (2Mb genomic region flanking *cortex*) for the presence of any miRNAs in *Bicyclus anynana*, a model species which enjoys a comprehensive annotation of miRNAs expressed throughout wing development (*20*). Two miRNAs, *mir-193* and *mir-2788,* were found, ∼5.2 kb apart, in the intergenic region between *cortex* and the adjacent protein-coding gene *poly(A)-specific ribonuclease* (*parn*) (Fig. 1C). Phylogenetic analyses showed that *mir-193* is deeply conserved across the animal kingdom, while *mir-2788* is conserved in Arthropoda (*21–23*) (Fig. S1).

To elucidate the functions of both miRNAs, we performed embryonic CRISPR-Cas9 mediated knock-out experiments in *B. anynana*. Single guide RNAs (sgRNAs) were designed to disrupt the essential Drosha or Dicer processing sites in the miRNA biogenesis pathway to effectively block the biogenesis of mature miRNAs (Fig. 1A-B, fig. S2, table S1)(*24*). F0 mosaic knock out (mKO) mutants exhibited similar phenotypes for the two miRNAs, but at different frequencies. For *mir-193*, 74-83% of the mKO mutants showed reduced melanin levels across all the body parts covered with scale cells, such as wings, antenna, legs, and abdomen (fig. S3, table S2). Variations in wing color were observed across individuals and sexes (fig. S3). Many of the mosaic mutants could not fly due to wing hinge weakness. For *mir-2788*, similar color phenotypes were observed, but only 0-8% of the adults showed wing color changes, and no flight defects were observed (fig. S4, table S2). Genotyping of mKO mutants, via next generation sequencing, suggested that various *mir-193* mKO phenotypes are likely the result of different, short, on-target mutant alleles, while the rare phenotypes observed in the *mir-2788* mKO mutants are likely the result of long, off-target adjacent disruptions, probably in *mir-193* (fig. S5-6, table S3, Supplementary text).

To draw a direct genotype-phenotype association, we crossed the F0 mKO mutants and generated homozygous mutant lines with short deletions for both miRNAs. (Fig. 1D, fig. S7). For *mir-193*, we generated four mutant lines with a 4bp (m5), 6bp (m4), 19bp (m2), and 24bp (m1) deletions around the *mir-193* 5’ Drosha processing site (Fig. 1D). A phenotypic series was observed where m1, m2, and m4, with 6-24bp deletions, showed almost equivalent light brown wing color, with “black disk” eyespot wing color patterns turning white (Fig. 1D, fig. S7). Homozygotes of these lines could not fly due to severe wing hinge weakness. Homozygotes of m5, with a shorter 4bp deletion, showed milder phenotypes. They had slightly darker wing colors, black “black disks”, and were able to fly (Fig. 1D, fig. S7). Heterozygotes of all the four mutant lines were visibly WT-like, in both color and behavior, indicating the recessive nature of these alleles. Quantification of the level of the guide strand of *mir-193*, miR-193-3p, using qPCR across mutants and WT pupal wings indicated that increasing levels of miR-193-3p correlated with darker wing colors across a phenotypic gradient, suggesting a dose-dependent effect of *mir-193* in wing melanization (Fig. 1E, table S4). For *mir-2788*, one mutant line was generated with a 14bp deletion around the 5’ Drosha processing site (Fig. 1D). No visible phenotypic changes were observed in both mutant heterozygotes and homozygotes, although both mature miRNA strands in the mutant line were depleted (Fig. 1D-E, fig. S7, table S4). This indicates that *mir-193*, but not *mir-2788*, promotes melanic wing color in *B. anynana*.

Four protein coding genes within the hotspot *cortex* locus, *cortex*, *parn*, *lemur tyrosine kinase* (*LMTK*), and *washout* (*wash*), were previously proposed to be potential color regulators in *Heliconius* butterflies (*15, 25, 26*). We asked whether these genes might also act as effectors within the locus. We generated mKO mutants of these genes using CRSPR-Cas9 in *B. anynana*, but none of the confirmed F0 mutants showed substantial changes in wing color (Fig. 1C, fig. S2, 8-10, table S1-2, 5, Supplementary text). This suggests that *mir-193* is the major effector within the locus, at least in *B. anynana,* a nymphalid butterfly.

To assess the functional conservation of *mir-193* beyond nymphalids, we tested its function in a pierid (*Pieris canidia*), and a papilionid (*Papilio polytes*), representatives of families where melanic polymorphisms were also previously mapped to the *cortex* locus (Fig. 1F, fig. S2, table S1). Reduced melanin pigmentation was observed in both species with high frequencies (55% - 58%) (Fig. 1G-H, fig. S11-12, table S2). In *P. canidia*, all the black/grey wing color patterns became white, while the yellow color remained unchanged (Fig. 1G, fig. S11). In *P. polytes*, the black wing color disappeared, exposing a variety of white/yellow/red colors across the wing (Fig. 1H, fig. S12). This suggests that the function of *mir-193* in promoting melanization is conserved across three major butterfly families that diverged around 90 My ago (*27*) (Fig. 1F).

## *Mir-193* is derived from, and is the functional unit of, a gigantic lncRNA *ivory*

Identifying the primary transcripts of miRNAs (pri-miRNAs) is essential to elucidate their transcriptional control and tissue/cell-specific expression patterns. Due to the high turnover rate of Drosha processing, pri-miRNAs are usually expressed at low levels and are not easily captured in RNA-seq data (*29*). In fact, none of the annotated lepidopteran genomes showed any annotated transcripts overlapping the two miRNAs in the *cortex* locus. Blocking Drosha processing, however, can trigger the accumulation of these primary transcripts (*29*). Taking advantage of the miRNA mutant lines that disrupted the Drosha processing sites, we performed RNA-seq to profile whole transcriptomes from sib-paired WT and mutant *mir-193* and *mir-2788* homozygotes. For *mir-193*, wing tissues from sib-paired female WT and m4 homozygotes were sequenced from 60% wanderer stage (late 5^th^ instar larval stage), Day 1 pupal stage, and Day 6 pupal stage. For *mir-2788*, wing tissues from sib-paired female WT and mutant homozygotes were sequenced from Day1 pupal stage alone. By inspecting all the RNA-seq data we found a gigantic transcript spanning a 370kb chromosomal region in two *mir-193* Day 1 and one *mir-193* Day 6 mutant pupal wing libraries, but not in the *mir-193* mutant larval wings, WT wings, or *mir-2788* mutant wing libraries (Fig. 2A). No open-reading frame was found in this newly discovered transcript, rendering it as a long non-coding RNA (lncRNA). There was no clear sequence homology for the lncRNA across Lepidoptera, except for the ∼100bp core promoter region surrounding its transcription start site (TSS) recovered by 5’ rapid amplification of cDNA ends (5’RACE) (fig. S13, table S6). This corresponds to the core promotor region of ‘*ivory*’, a lncRNA discovered in two recent preprints (*30, 31*). Thus, this newly annotated lncRNA is termed *ivory* thereafter.

**Fig. 2.**
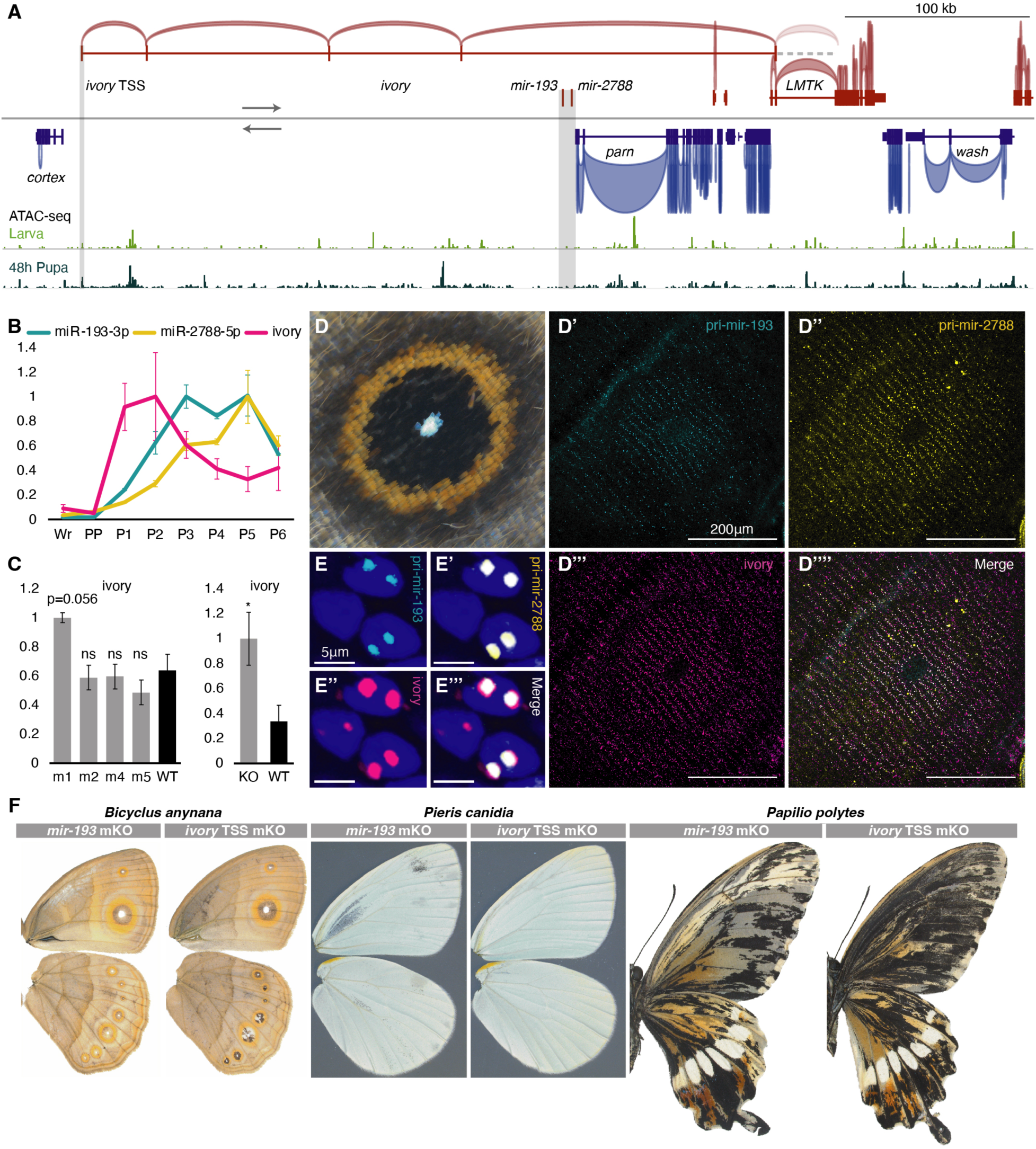
The lncRNA *ivory* functions as primary *mir-193*. **(A)** A gigantic lncRNA, *ivory*, was found in the RNA-seq data of three *mir-193* mutants, with a deeply conserved transcription start site (TSS) whose chromatin accessibility increased during the larval-pupal transition (shaded), and an undefined 3’ terminus. No open chromatin was found around the miRNA region (shaded). **(B)** Time series expression of miRNA mature strands, miR-193-3p, miR-2788-5p, and *ivory*. **(C)** Expression levels of *ivory* across the miRNA mutants and WT. n=3-4; ns: not significant; *: p<0.05; **: p<0.01; ***: p<0.001; Error bar: SEM. (**D-D’’’’**) Spatial expression (HCR) of *pri-mir-193*, *pri-mir-2788*, and *ivory* in the *B. anynana* ‘eyespot’ wing color pattern and (**E-E’’’**) their expression signals within individual nuclei. **(F)** *mir-193* and *ivory* TSS mosaic knock-out (mKO) phenotypes in *B. anynana*, *P. canidia*, and *P. polytes*. Images were horizontally flipped when necessary.

In *B. anynana*, one intron of *ivory* overlaps the two miRNAs and *ivory* is the sole transcript overlapping the two miRNAs with the same transcription orientation (Fig. 2A). Thus, we hypothesized that *ivory* is the primary transcript of *mir-193* and *mir-2788*. To test this hypothesis, we first generated a time series expression profile for the guide strands of the two miRNAs, as well as for *ivory* (table S4). As expected, the three non-coding RNAs exhibited very similar expression profiles – negligible expressions in larval wings and high expressions in pupal wings (Fig. 2B). Noticeably, *ivory* exhibited an expression peak earlier in development than the two mature miRNAs, which aligns with the general pattern of miRNA processing in *Drosophila*: a fast turnover rate of the primary transcript produces long-lasting mature miRNA products (*32*). Furthermore, the expression pattern of *ivory* during the larval-pupal transition correlated with an increasing chromatin accessibility around the *ivory* promotor, suggesting a causal relationship (Fig. 2A, fig. S13).

As miRNA primary transcripts are supposed to accumulate upon the disruption of the Drosha processing sites, we quantified the level of *ivory* across all *mir-193* and *mir-2788* mutant lines (table S4). Across the four *mir-193* mutant lines, the lncRNA is marginally (p=0.056) overexpressed in m1 mutants, compared with WT, but not in the other mutants with shorter deletions (Fig. 2C). In the *mir-2788* mutants, the lncRNA is significantly overexpressed compared with WT (Fig. 2C). This provided moderate evidence that the two miRNAs were derived from *ivory*.

Next, we used hybridization chain reaction (HCR) to examine the spatial expression of the primary transcript of the two miRNAs as well as *ivory* (Data S1). Expression signals of *pri-mir-193*, *pri-mir-2788*, and *ivory* mapped to the black/brown color of *B. anynana* wing patterns (Fig. 2D-D’’’’). The overlapping expression signals for the two pri-miRNAs and *ivory* appeared as two nuclear dots, corresponding to the two chromosomal transcription sites (*33*) (Fig. 2E-E’’’). Transcription of intronic miRNAs can potentially involve both host gene-dependent, and independent transcriptional regulations (*34*). However, there was insufficient evidence to support an alternative *mir-193* TSS independent of *ivory* (fig. S14, table S6, Supplementary text). To further validate whether *ivory* carries the same function as *mir-193*, the deeply conserved *ivory* TSS was disrupted using CRISPR-Cas9 in *B. anynana*, *P. canidia*, and *P. polytes* (fig. S2, table S1). The mKO mutants exhibit reduced melaninization at high frequencies (54%-98%), and the *ivory* TSS mKO mutants completely phenocopied the *mir-193* mKO mutants in each species (Fig. 2F, fig. S9, 15-17, table S2). These experiments show that the lncRNA *ivory* serves as the primary transcript for *mir-193/2788*, and that *mir-193* is the functional product of *ivory*.

## *Mir-193* directly target multiple pigmentation genes

MiRNAs elicit their regulatory effects by repressing target mRNAs. To discover both direct and indirect targets of *mir-193*, we examined the transcriptomes generated from the *B. anynana* m4 mutants and sib-paired WT. A total number of 4 (larva), 11 (Pupa day1), and 218 (Pupa day6) differentially expressed genes (DEGs, padj<0.01) across mutant and WT wings were found, suggesting that *mir-193* triggers large transcriptomic changes mostly during late pupal development, when it is highly expressed (Fig. 3A, Data S2). Except for an uncharacterized protein (NCBI gene ID: LOC112054702) ∼680kb away from *cortex*, none of the genes within the *cortex* locus (2Mb genomic region flanking *cortex*) appeared as DEGs, suggesting the *trans*-, rather than *cis*-acting nature of the miRNA. Multiple genes previously associated with ommochrome and melanin pigmentation pathways in butterfly wings appeared as DEGs in Day 6 pupal wings (*35*).

**Fig. 3.**
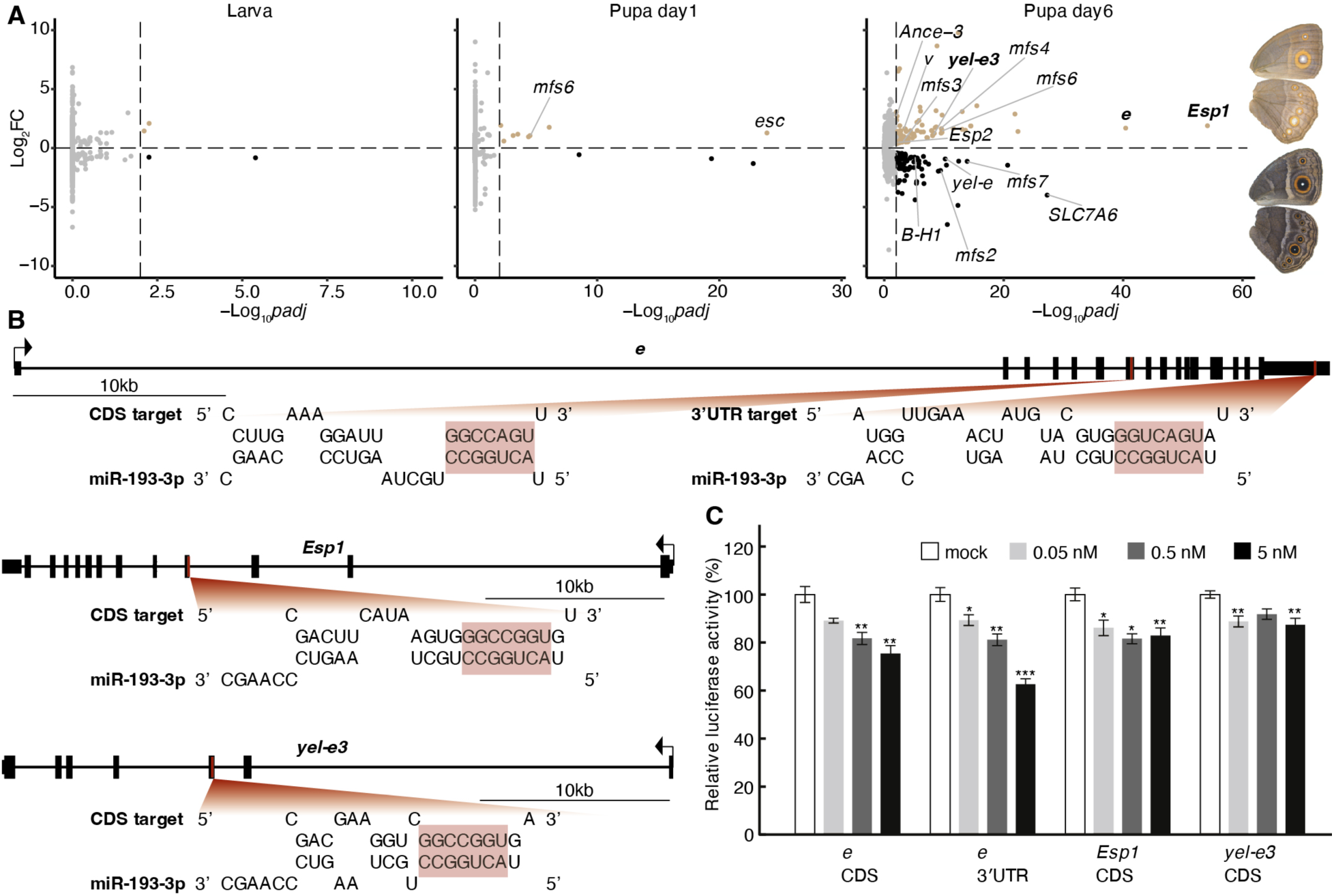
*mir-193* directly targets multiple pigmentation genes. **(A)** Differentially expressed genes (padj<0.01) across sib-paired female *mir-193* m4 mutant and WT wing tissues across wing development. Genes highlighted were previously associated with butterfly pigmentation, or potential color regulators. Candidates for in-vitro validation are in bold **(B)** Four putative binding sites of miR-193-3p, the guide strand of *mir-193*, were found in CDS and/or 3’UTR regions of three candidate genes, *ebony* (*e*), *Esp1*, and *yellow-e3* (*yel-e3*) that are upregulated in Day6 mutant wings. Full sequence complementarity (allowing G:U wobble base-pairing) between the seed region (nucleotides 2-8 from the 5’ terminus of the miRNA guide strand) and the miRNA targets are highlighted. **(C)** Dual luciferase reporter assay was used to validate the miRNA-target silencing across the four predicted binding sites in-vitro, with a concentration gradient of miR-193-3p mimic. n=6; *: p<0.05; **: p<0.01; ***: p<0.001; Error bar: SEM.

Since direct targets of *mir-193* are expected to be overexpressed in the miRNA mutants, all genes up-regulated in the mutants were pooled and searched for complementary binding sites to miR-193-3p, the guide strand of *mir-193*. In total, 49 out of 118 genes highly expressed in the mutant were predicted to be direct targets of miR-193-3p (Data S3). Three candidate targets, *ebony* (*e*), *Esp1*, and *yellow-e3* (*yel-e3*), were chosen for further investigation. *e* is a well-known insect melanin pathway gene that catalyzes the conversion of dopamine to NBAD – a light yellow pigment. *Esp1* is associated with ommochrome color patterns in *V. cardui* butterflies, and *yel-e3* is a melanin pathway gene, but both have unknown functions (*35*). Four putative miR-193-3p binding sites were found in either the protein coding sequence (CDS), or the 3’ untranslated region (3’UTR) of these genes (Fig. 3B).

Using a dual luciferase reporter assay, we validated the silencing effect of miR-193-3p across all the predicted target sites in-vitro (Fig. 3C, table S7, Supplementary text). Moreover, miR-193-3p exhibited a dose-dependent silencing effect on the two *e* target sites (Fig. 3C). Based on the known function of *e*, we propose that when *mir-193* was disrupted in the black/brown wing regions in the m4 mutant, *e* was de-repressed in those regions, and started converting dopamine to NBAD, producing a lighter color, and shunting dopamine away from melanin pigment production (*35, 36*).

## *Mir-193* is an ancestral melanic color regulator

Finally, to test whether regulation of melanic coloration, via *mir-193,* is ancestral to the Lepidoptera, we tested its function in *Drosophila melanogaster,* an outgroup species. In flies, *mir-193* is in a different genomic region, lacking *ivory*, and between two protein-coding genes with the same transcription orientation (Fig. 4A). To determine the function of *mir-193* in *D. melanogaster*, we used the *pannier* Gal 4 driver (*pnr-Gal4*) to express either a *mir-193* sponge to repress *mir-193* function, or extra *mir-193* precursors to enhance *mir-193* function. As a result, in the *mir-193* sponge line, female A6 segments became brighter, while in the *mir-193* overexpression line, female A6 segments became darker, compared with the control line with a similar genetic background (Fig. 4B). This suggest that *mir-193* has an ancestral role in promoting melanization in flies and lepidopterans, and regardless of genomic context (Fig. 4C).

**Fig. 4.**
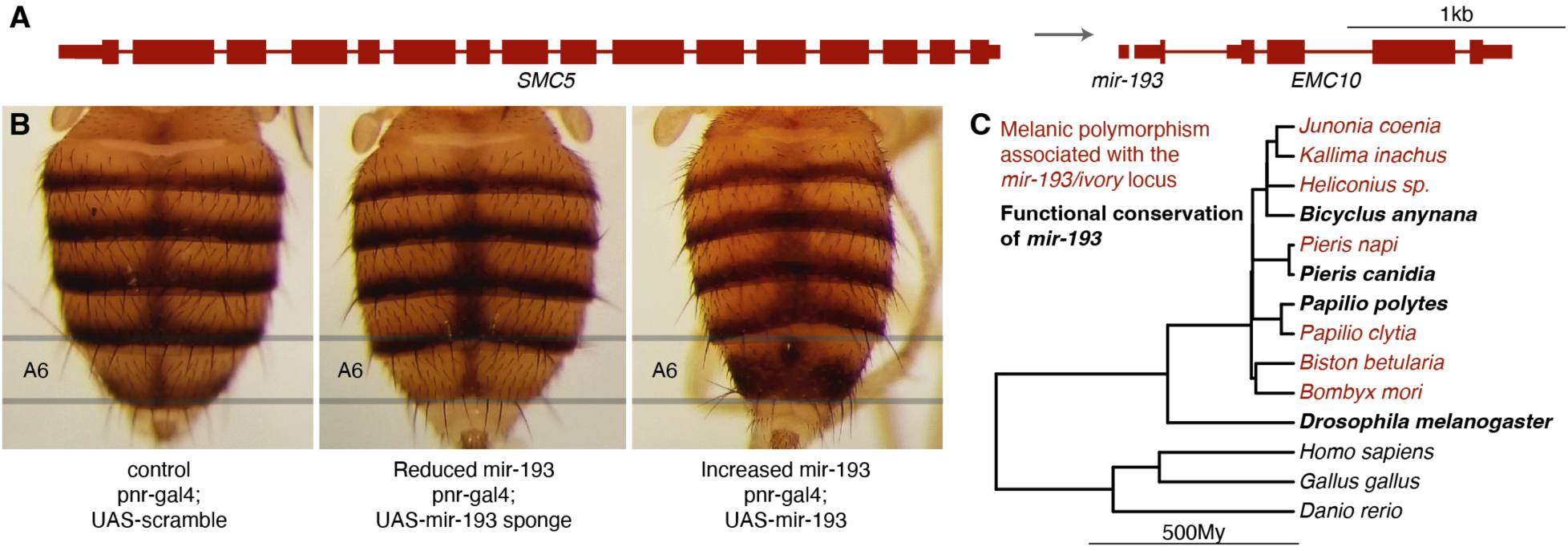
*mir-193* is an ancestral melanic color regulator. **(A)** Genomic context of *D. melanogaster mir-193*. **(B)** Phenotypes of *pannier* Gal4 driver (*pnr-Gal4*) transgenic lines expressing either a *mir-193* sponge with a sequence of seed-complementary binding sites for the guide strand miR-193-3p (reduced *mir-193*), or extra *mir-193* precursors (increased *mir-193*), or a *mir-193* sponge with scrambled miR-193-3p binding sites (control). **(C)** Functional conservation of *mir-193* across a broader animal phylogeny where *mir-193* is deeply conserved. Phylogeny is from (*28*).

Genotype-phenotype association studies are commonly used to assign variation in a genomic region to variation in a phenotypic trait. In many cases, the causative mutations are frequently found in intergenic regions, and it is usually assumed that these regions harbor cis-regulatory elements (CREs) that regulate the expression of flanking genes. When the effector gene is a poorly annotated non-coding element that acts in *trans*, such as *mir-193* or its primary transcript *ivory* investigated in this study, a direct link between local genetic variation and flanking protein-coding genes can lead to misleading conclusions. This happened with the *cortex* locus where a large number of intraspecific wing color pattern polymorphisms in lepidopterans, in most cases involving black and white, or dark and bright color patterns, have been mapped to this hotspot locus and been mistakenly attributed to *cortex* regulation (*2, 5, 7, 15, 17*). It will be interesting to examine in future how the genomic contexts and targets of *mir-193* have evolved across animals and how, *ebony*, one of the targets of *mir-193* in *B. anynana*, became the hotspot locus across *Drosophila* species (*37*), despite the presence and conserved function of *mir-193*.

Our study identified a micro-RNA, processed from a long non-coding RNA, as the likely effector of a hotspot locus that underlies adaptive evolution in animals. This adds to a recent discovery of small non-coding RNAs being key regulators of adaptive flower color evolution and speciation (*38*). The burst of miRNA innovation at the base of Lepidoptera (*21, 22*), may have served as evolutionary raw materials to create a splendid gamut of morphological diversity within this order, one of the most species-rich on earth. This and future investigations on non-coding RNAs will shed light on the long-standing hypothesis that it is the complexity of swiftly evolving non-coding components of the genome (cis-acting regulatory DNA elements and trans-acting non-coding RNAs), rather than the relatively static proteome, that drives organismal complexity (*39, 40*).

## Acknowledgments

We thank Dr. Shinya Komata and Dr. Nicholas Vankuren for their suggestions on rearing and performing CRISPR experiments on *P. polytes*. We thank members of the Facebook group ‘The Hungry Caterpillar Singapore’ for donating lime plants for rearing *P. polytes*. We thank prof. Nalini Puniamoorthy for providing additional lab space to host the *B. anynana* miRNA mutant lines. We thank prof. Yvonne Tay for the helpful comments on this work.

## Author contributions

Conceptualization: ST, AM

Methodology: ST, TDB, AM

Investigation: ST, YA, TDB, JLQW, AL, YW, SNM

Visualization: ST, YA, TDB, AL, YW

Funding acquisition: AM, PJW, KUT, YA

Project administration: AM, PJW, KUT

Supervision: AM, PJW, KUT

Writing – original draft: ST, AM

Writing – review & editing: ST, AM, PJW, KUT

## Data and materials availability

All data are available in the main text or the supplementary materials. Raw RNA-seq and ATAC-seq data will be uploaded onto the NCBI database.

## Supplementary Materials

Materials and Methods

Supplementary Text

Figs. S1 to S17

Tables S1 to S7

References (*1–15*)

Data S1 to S3

## Supplementary Materials

### Materials and Methods

#### Insect husbandry

Lab populations of *B. anynana, P. canidia,* and *P. polytes* butterflies were reared 27°C, with a 12:12 day: night cycle and 60% relative humidity. *B. anynana* larvae were fed young corn leaves and adults were fed mashed banana. *P. canidia* larvae were fed potted cabbage leaves and adults were fed artificial nectar. *P. polytes* larvae were fed lime leaves and adults were fed artificial nectar.

#### CRISPR-Cas9 genome editing

Embryonic CRISPR-Cas9 knock out experiments were performed in *B. anynana*, *P. canidia*, and *P. polytes* following the established protocol (with a video protocol) with some modifications (*1*). For miRNAs, guide RNAs were designed to target the Drosha or Dicer processing sites to disrupt mature miRNA biogenesis. For protein coding genes, guides were designed within the coding region of an exon shared by all isoforms of the target gene, and close to the 5’ termini to effectively induce reading frame shift in the coding region. For the lncRNA *ivory*, guides were designed on its highly conserved TSS to disrupt its transcription. One or two guides were designed for each target and each guide was used independently. Specificity of the guide sequences was checked by blasting against the genome of each species. Only those with a unique hit were selected.

For all the guides used in *B. anynana* (except for the two guides targeting *cortex*), template single guide DNAs (sgDNAs) were produced by PCR, using NEB Q5 High-Fidelity DNA Polymerase. sgDNAs were transcribed into sgRNAs using NEB T7 RNA Polymerase. sgRNAs were purified by ethanol precipitation. Size and integrity of sgDNAs and sgRNAs were checked using gel electrophoresis. For the two *B. anynana* guides targeting *cortex*, and all guides used in *P. canidia* and *P. polytes*, synthetic crRNAs were ordered from Integrated DNA Technologies (IDT).

For microinjection of sgRNAs in *B. anynana*, corn leaves were placed in adult cages around 2-3pm. Leaves were left in the cages for 1h and eggs were collected immediately and placed onto 1mm-wide double-sided strips of sticky tape in petri dishes. We used a final concentration of 500ng/μL IDT Alt-R S.p. Cas9 Nuclease V3 (with 1 x Cas9 buffer), and 250ng/μL sgRNA in the injection mixture. The injection mixture was incubated at 37 °C for 10 min prior to injection. 0.5 μL food dye was added per 10 μL mixture to facilitate the visualization of the injected solutions. Microinjection was performed using glass capillary needles. Wet cotton was placed inside the petri dishes with injected eggs, and the eggs were reared at 27°C (wet season (WS) condition) in most cases, except for *mir-2788*, when some batches of injected eggs were reared at 17 °C (dry season (DS) condition). Newly hatched larvae were feed cut corn leaves in small containers, then shifted to corn plants in cages to complete development.

Microinjection of synthetic crRNAs for the two *cortex* guides in *B. anynana* and all guides in *P. canidia* and *P. polytes* followed the same guide preparation steps but with different egg collection strategies. *B. anynana* eggs were collected as described before. *P. canidia* eggs were collected using a small plastic container lined with a piece of parafilm and topped with a fresh cabbage leaf. This setup encouraged adult butterflies to deposit most of their eggs onto the parafilm, enhancing hatchlings’ survival rates by preventing egg dehydration. The container was left inside the cage for approximately 4 h to optimize the number of eggs collected. Once a sufficient number of eggs were obtained, both the leaf and the parafilm were transferred to a Petri dish for injection. *P. polytes* eggs were collected using fresh lime tree cuttings. Lime tree cuttings with young leaves were placed in the adult cages with hand-paired females for approximately 1-2h to optimize the number of eggs collected. Eggs were collected immediately and placed onto 1mm-wide double-sided strips of sticky tape in petri dishes, with the bottom of each egg with a flat surface facing up to facilitate injection. For all three species, eggs were injected with CRISPR/Cas9 protein-crRNA-tracrRNA complexes. Stock solutions of crRNAs and tracRNA were prepared as 1000ng/μL, 4μL of crRNA was incubated with 4μL of tracrRNA and 12μL of Nuclease-Free Duplex Buffer at 95°C for 5 mins and then left to cool down at room temperature. We used a final concentration of 500ng/μL Cas9 protein (with 1 x Cas9 buffer), and ∼ 200ng/μL annealed crRNA-tracrRNA in the injection mixture. After injection, moistened cotton balls were then placed into each Petri dish. Hatchlings were promptly transferred to the host plants of the corresponding species and raised under the WS condition until they reached adulthood. All newly emerged butterflies were frozen at −20 °C ∼6h after adult emergence. Phenotypic changes were inspected manually and mutants were imaged using a Leica DMS 1000 microscope. All the guide sequences were listed in table S1. All the CRISPR injection statistics were summarized in table S2.

#### In-vivo T7 endonuclease assay

In-vivo T7 endonuclease assays were performed to confirm the cutting efficiencies of all guides used in *B. anynana* and *P. polytes*. Genomic DNA was extracted from ∼20 injected eggs 4 days after injection, using E.Z.N.A Tissue DNA Kit. For each miRNA, one pair of CRISPR genotyping (GT) primers was designed to amplify a 300∼600bp genomic region spanning the target sites using PCRBIO Taq Mix Red. PCR products were purified using ThermoFisher GeneJET PCR Purification Kit. 200ng of PCR products were denatured and re-hybridized with NEB Buffer 2 in a total volume of 20μL following the reported temperature settings (*1*). The product was divided into two tubes, one treated with 1μL T7 endonuclease (New England Biolabs), and the other without endonuclease as a control. Both were incubated at 37°C for 15min and were subsequently run on a 2% agarose gel. All guides showed efficient cutting of the designated target sites in-vivo, except one *parn* guide in *B. anynana*, due to single nucleotide polymorphisms (SNPs) present in the guide region (fig. S2).

#### Genotyping mosaic mutants via Sanger sequencing

On-target mutations across mKO mutants (except for the mKO mutants of the two miRNAs) in *B. anynana*, regardless of the presence of any phenotypic changes, were checked using Sanger sequencing. Genomic DNA was extracted from thorax of the mKO mutants using E.Z.N.A Tissue DNA Kit. The GT primers used in the T7 assay were used to amplify the genomic region spanning the target sites using PCRBIO Taq Mix Red. PCR products were purified using ThermoFisher GeneJET PCR Purification Kit, and were sent for Sanger sequencing by 1^st^ Base (Axil Scientific Pte Ltd, Singapore). Guide-induced indels were checked using the Synthego ICE Analysis tool v3. Genotyped individuals and their indel rates were shown in table S5. Examples of the mKO phenotypes and their corresponding genotyping results were shown in fig. S9.

#### Genotyping mosaic mutants of the miRNAs via NGS

F0 mKO mutants of the two miRNAs in *B. anynana* were genotyped via next generation sequencing (NGS). For *mir-193*, genomic DNA was extracted from five mosaic mutant hindwings with homogeneous but different melanin levels, from four male mKO mutants (fig. S5). For *mir-2788*, genomic DNA was extracted from both hindwings of a male mosaic mutant with paired phenotypes, one wing showed reduced melanin levels, while the other wing appeared WT-like (fig. S6). DNA extraction was performed using E.Z.N.A Tissue DNA Kit. For *mir-193*, a 362bp region around the sgRNA cutting site was amplified from genomic DNA with Illumina genotyping primers, while for *mir-2788*, a 3820bp region was amplified with Pacbio genotyping primers. PCR was performed using NEB Q5 High-Fidelity DNA Polymerase and PCR products were purified using ThermoFisher GeneJET PCR Purification Kit. PCR products were checked using gel electrophoresis and NanoDrop, and were sent for NGS. The Illumina and Pacbio genotyping primers were listed in table S3.

For *mir-193*, Illumina NovaSeq short-read amplicon sequencing was performed for the five mutant wing samples to generate 1M 250bp paired-end reads for five samples in total. For *mir-2788*, Pacbio Sequel II long-read amplicon sequencing was performed for the two wing samples to generate 7K-10K 4kb HiFi reads per sample. Quality check, library construction, and sequencing were all carried out by Azenta, Singapore. For Illumina short reads, adaptors were trimmed using Cutadapt. Clean reads were aligned to the reference sequence of the amplified region, and mutant alleles were called using CRISPResso2. Pacbio long reads were aligned to the reference sequence of the amplified region using NGMLR, and alignments were visualized in Geneious. Structural variants and mutant alleles were called using Cas-Analyzer and Sniffles.

#### Generation of miRNA mutant lines

Homozygous mutant lines were generated for both *mir-2788* and *mir-193* in *B. anynana*. In general, F0 mosaic female mutants with wing phenotypes from CRISPR knock out experiments were crossed with WT males. Some of these F0 mosaic females might carry one or several mutant alleles in their germline. F1 offspring were screened via hemolymph extraction and Sanger sequencing, as described below. F1 heterozygotes carrying mutant alleles were crossed with either other F1 heterozygotes carrying the same mutant allele, if any, or with WT individuals. Subsequent generations were screened in the same way, and crossed until mutant homozygous lines were generated. The *mir-2788* mutant line was maintained in a completely homozygous status. For *mir-193*, the four mutant lines were maintained in a way that in each generation, mutant homozygous females (with visible wing phenotypes) were crossed with heterozygous males (WT-like), since *mir-193* mutant alleles were recessive and homozygous mutants have various degrees of wing hinge weakness. WT-like heterozygous males were needed to perform proper courtship displays for successful mating.

#### Genotyping mutant lines via hemolymph extraction and Sanger sequencing

For hemolymph extraction, around 10μL hemolymph was extracted from 5^th^ instar larvae and suspended in 200μL Saline-Sodium Citrate buffer. Hemocytes were collected by centrifugation, then resuspended and incubated in 20μL digestion buffer (1.1mL 1M Tris-HCl of pH6.8 in 50mL deionized water) with 2μL proteinase K (New England Biolabs) at 37°C for 15min. Upon heat inactivation of proteinase K at 95°C for 3min, 3μL of the cell lysate containing genomic DNA was used for PCR using the same GT primers from the CRISPR knock out experiments. PCR was performed using PCRBIO Taq Mix Red. PCR products were purified using ThermoFisher GeneJET PCR Purification Kit, and were sent for Sanger sequencing by 1^st^ Base (Axil Scientific Pte Ltd, Singapore). Zygosity of the amplicons were checked using the Synthego ICE Analysis tool v3.

#### qPCR

To quantify the expression of miRNAs and the lncRNA, all wings were sampled from females for precise staging and avoid dealing with variation between sexes. Staging of females was performed as described before. For *mir-193* and *mir-2788* mutants, wings were sampled at Day1 (15%) pupal stage, from both homozygous mutants and WT individuals. To generate a time series expression profile for the guide strand of the two miRNAs, miR-193-3p and miR-2788-5p, as well as *ivory*, wings were sampled from WT animals at 60% wanderer stage, 50% pre-pupal stage, and Day 1-6 (15% - 92%) pupal stages. One forewing and one hindwing from the same individual were dissected and pooled as one biological replicate. Each condition consists of 3-4 biological replicates. Fresh tissues were preserved in RNAlater and stored at −80 °C before RNA extraction.

Total RNAs, including small RNAs, were extracted using miRNeasy Tissue/Cells Advanced Mini Kit. To quantify mature miRNAs, cDNAs were reverse-transcribed from 1μg total RNAs using stem-loop (SL) primers designed for each miRNA, together with an SL primer for the small nuclear RNA (snRNA) U6 as an internal control (*2*). To quantify *ivory*, cDNAs were reverse-transcribed from 400ng total RNAs using oligo-dT. *Rps18* was used as an internal control. Reverse transcription was performed using RevertAid RT Reverse Transcription Kit. To quantify mature miRNAs, cDNA products from the reverse transcription reaction were used for qPCR using miRNA-specific qPCR forward primers, and a universal qPCR reverse primer. To quantify *ivory*, qPCR primers for *ivory* were used. qPCR was performed using KAPA SYBR FAST Universal Kit. The qPCR thermal cycling program was set as 40 cycles of 95°C for 10s and 60°C for 30s. For each biological replicate, three technical replicates were included, and expression data was analyzed using the 2^-ΛΛCt^ method. Standard curves were generated for qPCR primer pairs to ensure high primer efficiencies. Melting curves were also generated after each qPCR run and checked to ensure unique amplification. All SL primers and qPCR primers were listed in table S4.

#### In-situ HCR

Detection of *ivory*, *pri-mir-193*, and *pri-mir-2788* in Day1 pupal wings of *B. anynana* was carried out either automatically using an inbuilt robotic fluidics rig or manually based on the protocol described in (*3*) with few modifications in buffers and incubation conditions. For *ivory*, probes were designed on the first exon. For the two pri-miRNAs, probes were designed on the 1kb genomic region flanking the miRNA precursors, as described before (*4*). Briefly, wings were dissected in 1x PBS and transferred to glass chambers where they were fixed in 1x PBST supplemented with 4% formaldehyde for 30 mins. After fixation, the tissues were treated with a detergent solution (*5*) and washed first with 1x PBST followed by washes in 5x SSCT. Afterward, wings were either mounted on a slide with 20µL of poly-lysine (Sigma-Aldrich) coating on top of each wing for the automatic procedure or the tissues were transferred to glass wells with 500µL of 30% probe hybridization buffer for the manual procedure. For both the manual and the automatic rounds, the hybridization involved incubation in a solution containing 50μL (50μM) of probe set against each gene (IDT) in 2000 µL of 30% probe hybridization buffer followed by rigorous washing with 30% probe wash buffer. Afterward, wings were washed with 5X SSCT and incubated in an Amplification buffer for 30 mins. For the chain reaction, a solution with HCR hairpins (Molecular instruments) in amplification buffer was added to the tissues followed by washes in 5x SSCT. Finally, the wings and embryos were mounted on an inhouse mounting media and imaged under an Olympus fv3000 confocal microscope. For the manual methodology, the primary incubation was carried out for 16 hrs, and the secondary hairpin incubation for 8 hrs. The automatic in-situ protocol was carried out within 10hrs using customized buffers at 37°C in a 3D printed chamber. All the HCR probe sets were listed in Data S1.

#### RNA-seq

RNA-seq was performed using wing tissues from sib-paired female *mir-193* m4 mutants and WT animals, and from sib-paired female *mir-2788* KO mutants and WT animals. For *mir-193*, wing tissues were sampled from 60% wanderer stage (late 5^th^ instar larval stage), Day1 pupal stage, and Day6 pupal stage. For *mir-2788*, wing tissues were sampled from Day1 pupal stage. One forewing and one hindwing from one female were used as one biological replicate, four replicates were included in each condition except for Day6 WT samples (coupled with *mir-193* mutants) and Day1 WT samples (coupled with *mir-2788* mutants), which involved 3 replicates. Fresh tissues were dissected and stored in RNAlater before total RNA extraction using miRNeasy Tissue/Cells Advanced Mini Kit. Poly-A enriched stranded mRNA sequencing libraries were constructed, over 30 million 150bp read pairs per sample were generated using Illumina Novaseq platform. Quality check, library construction, and sequencing were performed by Novogene, Singapore (for *mir-193* samples), or by Azenta, Singapore (for *mir-2788* samples).

Trimmomatic 0.39 was used to trim adaptors from raw sequencing reads (options: PE ILLUMINACLIP:TruSeq3-PE-2.fa:2:30:10:8:true MAXINFO:40:0.2 MINLEN:32). Adaptor-trimmed clean reads were used to quantify expressions of genes annotated in the NCBI *B. anynana* ilBic genome (*6*) using Salmon 1.6.0, with the quasi-mapping mode (options: --validateMappings --seqBias –gcBias). Gene expression levels were normalized, and differential gene expression analysis was performed using DESeq2 in R studio. All the defferentially expressed genes between WT and *mir-193* m4 mutants were listed in Data S2.

#### 5’RACE and Nanopore sequencing

5’ rapid amplification of cDNA ends (5’RACE) was performed to 1) extend the 1^st^-2^nd^ exons of *ivory* towards the 5’ side to annotate the transcription start site (TSS) of *ivory*: 2) directly extend the *mir-193* precursor towards the 5’ side to detect if there exist any potential alternative TSSs of *mir-193* independent of *ivory*. 5’RACE System for Rapid Amplification of cDNA Ends v2.0 (Invitrogen) was used following the manufacturer’s instructions. In brief, the first gene-specific primer 1 (GSP1) was used to generate cDNAs from 1μg total RNAs. cDNAs were purified using S. N. A. P column, and oligo-dC tail was added using terminal deoxynucleotidyl transferase (TdT). Two round of PCR were performed to amplify the dC-tailed cDNAs. In the first round, the dC-tailed cDNA was amplified using a nested GSP2 and the Abridged Anchor Primer. The primary PCR product from the first round of PCR was 1:100 diluted, and used as a template for a second round of nested amplification, using a nested GSP3 and the Abridged Universal Amplification Primer. The final product was purified using ThermoFisher GeneJET PCR Purification Kit and sequenced using an Oxford MinION Flongle flow cell. 50ng of PCR product was used for library preparation with Rapid barcoding kit 24 V14 SQK-RBK114.24, following the manufacturer’s protocol. Each sample was individually barcoded before being pooled together for sequencing. The sequencing was performed on a Flongle Flow Cell (FLO-FLG114) with the r10 version and was run for 12 hours. Dorado 7.2.11 software was used for base calling. Reads were visualised in Geneious, and TSS was defined between the first nucleotide and the oligo-dC signal introduced in the 5’RACE experiment. All the GSPs were listed in table S6.

#### Annotation of *ivory* in *B. anynana*

For the discovery of unannotated transcripts in the *cortex* locus, all the adaptor-trimmed clean RNA-seq reads were mapped to the NCBI *Bicyclus anynana* ilBic genome using HISAT2 v2.2.1 and visualized in Integrative Genomics Viewer (IGV). The *ivory* transcript was annotated using both the aligned reads from three *mir-193* m4 mutant pupal wing libraries where *ivory* was captured, as well as the precise annotation of the *ivory* TSS using 5’RACE. Annotation of *ivory* at the 3’ end was up to the exon before it shares a common exon with *LMTK*, thus the annotated *ivory* transcript has an undefined 3’ termini. Sequence of the annotated *ivory* transcript (up to the exon before it overlaps *LMTK*) was shown in Supplementary text.

#### ATAC-seq

ATAC-Seq for pupal 24hr and 48hr forewing were performed as described in (*7*). We used 10 individuals and approximately 80,000 nuclei for each replicate, 3-4 replicates were involved in each condition. Forewings were collected and snap frozen in liquid nitrogen and stored in −80°C before nuclei extraction. Prepared libraries were sequenced using Novoseq 6000, with an average read depth of 30 million reads per library and 2*50 bp paired-end at Novogene, Singapore. 5^th^ instar larval forewing and pupal 3hr forewing ATAC-Seq data were from a previous publication (*7*) and downloaded from NCBI Bioproject PRJNA685019.

ATAC-seq analysis were carried out as described before (*7*). Reads obtained from the sequencing were processed using bbduk scripts from bbmap tools to remove any adapters. Reads were mapped to the NCBI *Bicyclus anynana* ilBic genome (*6*) using bowtie2. Reads mapped to the mitochondrial genome and marked as duplicates were removed using samtools idxstats and GATK MarkDuplicates respectively. Peak calling was done using *MACS2* (*8*) callpeak using --q 0.05 -- broad -f BAMPE -B --trackline as settings. The bedgraph output from the *MACS2* output were sorted to the chromosome coordinates and converted to bigwig file using *bedGraphToBigWig* and visualized in IGV.

#### miRNA target prediction

To find the direct targets of *mir-193*, all genes up-regulated in the *mir-193* m4 mutants across all three developmental stages were pooled. All annotated transcripts of the pooled genes were searched for their putative complementary binding sites of miR-193-3p, the guide strand of *mir-193*. In-silico miRNA target prediction was performed using RNAhybrid (*9*). The energy threshold was set to −20, with miRNA-target helix constraint in the seed region (nucleotide 2-8 from the 5’ terminus), allowing G:U wobble base-pairing in the seed. All predicted direct targets of *mir-193* were listed in Data S3.

#### Dual luciferase reporter assay

Dual luciferase reporter assay was performed to validate the silencing effect of miR-193-3p on the four predicted target sites in *e*, *Esp1*, and *yel-e3* in-vitro. An around 120-450bp 3’UTR or CDS region surrounding the predicted 3’UTR or CDS binding sites in each gene was used as a target. Target sequences were amplified from *B. anynana* genomic DNA and ligated into psiCHECK-2 vectors (Promega). For the 3’UTR target, e_3’UTR, the target was inserted into the 3’UTR region of renilla luciferase, for the three CDS targets, e_CDS, Esp1_CDS, and yel-e3_CDS, each of the targets was inserted just before the stop codon of the renilla luciferase CDS to create a renilla luciferase-target fusion protein construct. Double stranded miR-193-3p mimic was purchased from Ajinomoto Bio-Pharma, Japan. Hela cells were plated at a density of 2.0 x 10^5 cells/mL (0.5mL) in 24-well plates. After 24 hours, the medium was replaced with OPTIMEM, and the miR-193-3p mimic and each target vector were co-transfected using Lipofectamine 2000 (Thermo Fisher). Four hours after transfection, the medium was replaced with DMEM containing 10% FBS. Twenty-four hours after transfection, cells were collected and bioluminescence was measured using GloMax (Promega). The bioluminescent activity of the renilla luciferase was normalised to that of the firefly luciferase, the internal control. For each target construct, a concentration gradient of miRNA mimic was used (0.05nM, 0.5nM, and 5nM), with mock treatment without the miRNA mimic as a control. Each treatment involved six replicates. Sequences of miR-193-3p mimic and the tested targets were listed in table S7.

#### *mir-193* overexpression and sponge lines of *Drosophila melanogaster*

To determine whether fly *mir-193* had any effects on melanization in *Drosophila melanogaster*, we used the *pannier* Gal 4 driver (*pnr-Gal4*) to manipulate the abundance or availability of *mir-193.* This Gal4 driver activates the expression of coding sequences under the control of a UAS sequence along the dorsal abdominal mid-line during the pupal stages when adult pigmentation is developing and in females has the strongest expression in the A6 abdominal segment (*10, 11*). To reduce the number of copies of *mir-193* available to regulate expression of its target genes, we used competitive inhibition by crossing flies carrying *pnr-Gal4* to flies carrying the *UAS-mir-193-sponge* construct developed by (*12*), which carries 20 copies of a sequence complementary to the guide strand of *mir-193*, miR-193-3p, in the 3’ UTR of the reporter gene (Bloomington stock number 61398, genotype: w[*]; P{y[+t7.7] w[+mC]=UAS-mCherry.mir-193.sponge.V2}attP40; P{y[+t7.7] w[+mC]=UAS-mCherry.mir-193.sponge.V2}attP2). To determine the effects of increasing the abundance of *mir-193*, we crossed flies carrying *pnr-Gal4* to flies carrying a UAS-mir-193 construct (Bloomington stock number 59886, genotype: w[*]; P{y[+t7.7] w[+mC]=UASp-mir-193.S}attP16/CyO). Finally, to generate control flies with as similar a genetic background as possible, we crossed *pnr-Gal4* to a control stock made in parallel with the mir-193-sponge line but with a scrambled sequence in place of a *mir-193* binding site (Bloomington stock number 61501, genotype: w[*]; P{y[+t7.7] w[+mC]=UAS-mCherry.scramble.sponge}attP40; P{y[+t7.7] w[+mC]=UAS-mCherry.scramble.sponge}attP2). These control flies are expected to have wild-type levels of *mir-193*. All flies were aged 3 to 5 days after eclosion before imaging, and images were taken using a Leica MZ6 stereomicroscope equipped with a ring light and Scion (CFW-1308C) camera operated via TWAIN driver in Adobe Photoshop.

#### Statistical analysis

Effects of *mir-193* mutant alleles on the expression level of miR-193-3p (Fig. 1E, upper panel) was assessed using a generalized linear model in R.studio, using the glht function in the multicomp package. Tukey’s multiple comparison test was used to detect the differences in the expression levels of miR-193-3p across the *mir-193* mutants and WT. p<0.05 was considered significant. Two-tailed Student’s t-test was used in Excel to test the effect of *mir-2788* knock-out on the expression levels of the mature strands of *mir-2788* (Fig. 1E, lower panel), effect of miRNA knock-out on the expression levels of *ivory* (Fig. 2C), and the effect of miR-193-3p mimic treatments on the relative luciferase activity (Fig. 3C). p<0.05 was considered significant.

### Supplementary Text

#### Genotyping mKO mutants of *mir-193* and *mir-2788* in *B. anynana* via NGS

For *mir-193*, we genotyped five male mutant hindwings with homogeneous color changes, but with variable melanin levels in the eyespot black disk region (fig. S5). We used Illumina short-read amplicon sequencing to sequence a 362bp region around the sgRNA cut sites. We discovered that the mutation rate approached 100% in all the samples, and each had 1-3 major mutant alleles with indels of 3 to 123 bp in length (fig. S5). For *mir-2788*, we suspected rare long indels around the sgRNA cutting site that disrupted adjacent genomic causative regions. To capture these putative long indels, we performed Pacbio long-read sequencing of a 3820 bp region around the cut site from both hindwings of a male mutant, where one wing showed reduced melanin levels and the other wing had no visible phenotypic changes (fig. S6). Long deletions were detected in both wings only by one of the two analyses tools, but were not observed when the sequence alignments were inspected manually. However, short indels were observed at a near 100% rate around the cutting site in both samples (fig. S6). As a result, all the DNA extracted from the crispant wings from both *mir-193* and *mir-2788* knock-out experiments was near completely mutated with confirmed on-target short indels around the designated cutting sites. For *mir-193*, the various mutant phenotypes are likely the result of combinatorial effects of different mutant alleles around the cutting sites. For *mir-2788*, complete mutations with short indels around the miRNA locus were observed regardless of the presence of wing phenotypes, suggesting that the rare color phenotypes are not caused by the disruption of the small miRNA locus itself, but of a flanking genomic locus, potentially *mir-193*. In this case, the rare long deletion would extend beyond the 3.8kb amplicon as *mir-193* and *mir-2788* are ∼5.2kb apart, thus not captured in the Pacbio sequencing.

#### Test the roles of candidate protein-coding genes within the *cortex* locus

None of the mKO mutants of *cortex*, *LMTK*, and *wash* exhibited any visible phenotypic changes, while 19% of the mKO mutants of *parn* exhibited minor melanin reduction on the wing that was almost invisible (fig. S8). On-target mutations were confirmed across all the mKO mutants without any phenotypic changes, including *parn* (fig. S9). Without the generation of a homozygous mutant line, there’s insufficient evidence that *parn* regulates melanic color (fig. S8). In the case of *cortex*, by retrieving the previous RNA-seq data, we found that its expression was almost indetectable throughout wing development (TPM<1), further ruling out its function in wing coloration (fig. S10).

#### Test the alternative *mir-193* TSS using 5’RACE

To test if *mir-193* has an alternative TSS just upstream of the miRNA precursor, independent of *ivory*, we did 5’ RACE directly extending 5’ ends of the miRNA precursor. This recovered an alternative TSS 200 bp 5’ of the *mir-193* precursor only in one out of four wing tissues examined (fig. S14). This putative TSS, however, did not correlate with any open chromatin status across wing development, suggesting no genuine transcription signal (fig. S14). Moreover, no sequence homology of this site was found in any other Lepidoptera species. This provide insufficient evidence to support an alternative TSS of *mir-193*. The deeply conserved *ivory* TSS is likely the sole TSS of *mir-193*.

#### *ivory* sequence annotated in *B. anynana* (5’-3’, up to the exon before *ivory* overlaps *LMTK*)

ATTCGTCGTAGAGACCGCGAGCGGCGCGCGCGCGTTTCAACGAACTACGATTAAAAAAACAGTTCATTCAGCCGCCTCGGGAGCAACGACCTCGCGGCCATTATTTTTTCAACGAACGTCCTTGTCGCGAACAGCTGATTCACGAATCCGTGCTCCGAATGCAAAGTGATTCCTTGACAGAATGGAAAATAAGGTTTTTTAATCTCGTAGGTAATTTTCGATTGATACATTTTACAGTATAAAAATGTAGTGAGTCTATGTAGAACTATTTATTCTATATAACAATAAGGAATAGAGGAGTAACTGAAATTTGCAAAATTTGTTAATTATGACGTATTAGCGCTGAAATTTGTAAGCCCAATTTTAGTGTAGTGTAGTGTGGTAATTTTTCAATTTCAAAATGTGAAGTGTTTGTGTAAGTGATCGACATTGATCGGCAATCATCCAGGAACATTTCGGAATTTTTCGAGAATCTGCCGGACTAGTGTTTCGAGTAACCTTCAAAGCCAAAGATCTGCCTACACTATATTTCGGATTTATTTCGACCAGTTGTGTGGTGCCTAGTGAAACTATCGCCACAATTTATACAAAATACAGATGCTAATGGGTCAGTATTGGAGGCATGGAGCACCGAGACGGGGCTGTATGCAGTGTTGCCAGCGCTGGCGCTGGTCCTCACCGCTGTGGGCCTGCTGTTTGGCTGCACCTGGTGTTACAGGCACAAGGATTGTAAG

#### Validate the silencing effects of miR-193-3p on the predicted targets

Four putative miRNA target sites among three genes, *e*, *Esp1*, and *yel-e3*, were validated using dual luciferase reporter assay. Among the four target sites, one was within 3’UTR (*e*_3’UTR), and the rest were within protein coding regions (*e*_CDS, *Esp1*_CDS, *yel-e3*_CDS) (Fig. 3B). In fact, miRNAs are usually more effective in targeting 3’UTR than CDS (*13*). This aligned well with the fact that *e*_3’UTR showed relatively strong silencing effect compared with other targets (Fig. 3C). Moreover, weak G:U wobble base-pairs were present in the seed of *e*_3’UTR, *Esp1*_CDS, and *yel-e3*_CDS (Fig. 3B). Although the presence of G:U in the seed would abolish miRNA-mediated silencing in the case of seed-only complementarity (*13*), complementarity in a sub-seed region (nucleotides 13-16 from the 5’ terminus of the miRNA guide strand) would facilitate silencing (*14*). This aligned with the fact that although G:U pairs were present in the seed of *e*_3’UTR, *Esp1*_CDS and *yel-e3*_CDS, sub-seed complementarities were present in *e*_CDS, *e*_3’UTR, and *Esp1*_CDS (Fig. 3B), so some silencing effects can also be seen (Fig. 3C). To sum up, although significant silencing effects were validated experimentally among all four target sites, the most effective targets would be *e*_CDS or *e*_3’UTR, and conversely, the least would be *yel-e3*_CDS. The detailed miRNA-mediated silencing machineries aligned well with the assay data (Fig. 3B-C).

**Fig. S1.**
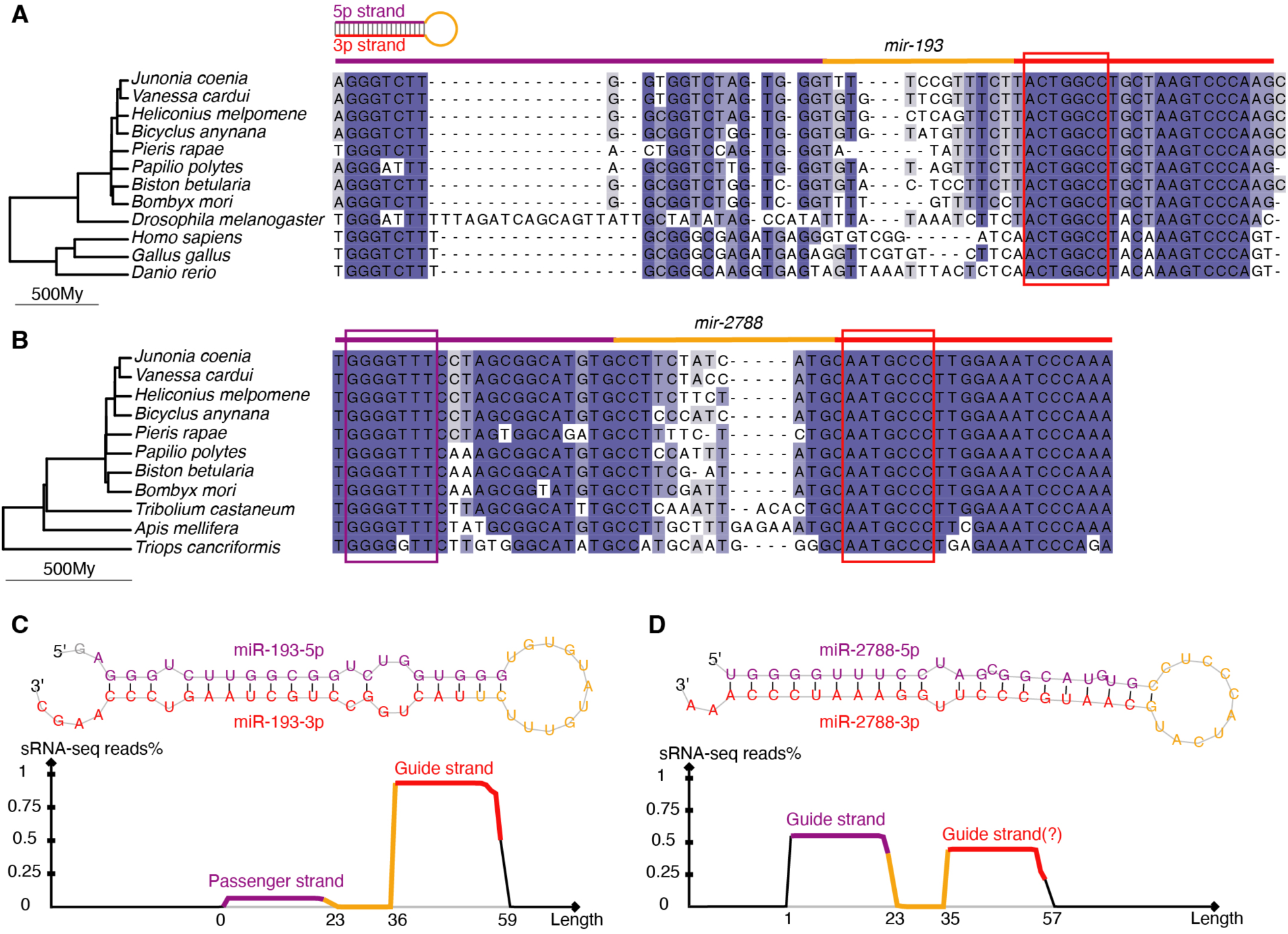
Sequence features of the two deeply conserved miRNAs embedded in the *cortex* locus. Deep sequence conservation of **(A)** *mir-193*, and **(B)** *mir-2788*, across the animal kingdom and across Arthropoda, respectively. Seed regions (nucleotides 2-8 from the 5’ terminus of the guide strand) that are essential for target recognition were squared. The structures of hairpin precursors of *mir-193* and *mir-2788* and the expression profiles of their mature miRNA strands are shown in **(C)** and **(D)**, respectively. Guide strand of a miRNA exhibits a substantially higher expression level than that of the passenger strand. When both strands are expressed at high levels (in the case of *mir-2788*), both can be potential guides. The expression profiles are from a previous small RNA-seq (sRNA-seq) dataset generated across wing development in *B. anynana* (15).

**Fig. S2.**
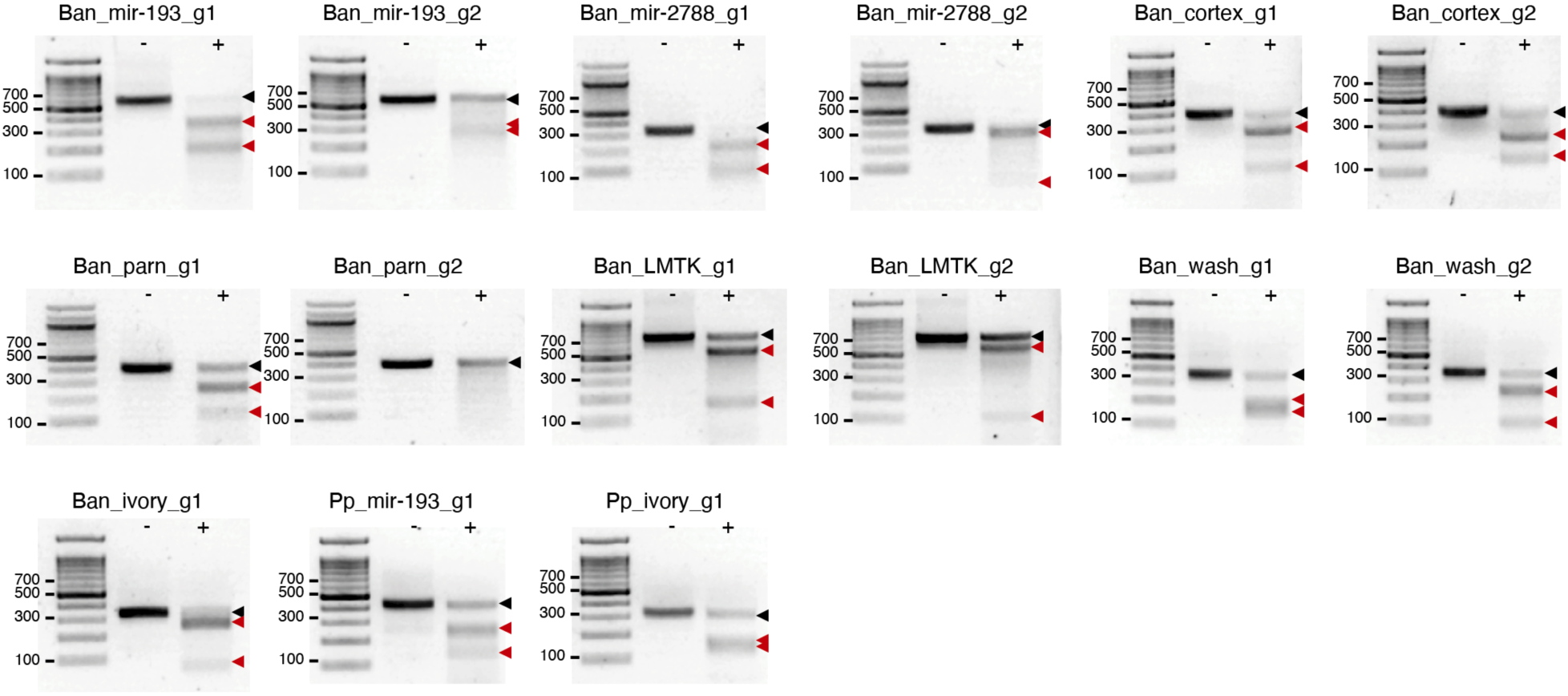
In-vivo T7 endonuclease assay. In-vivo T7 endonuclease assay was performed to test the cutting efficiencies of all guides used in *B. anynana* (Ban) and *P. polytes* (Pp). Control wells without T7 endonuclease treatments are denoted ‘-‘, while treatment wells are denoted ‘+’. The uncut band is indicated by a black arrowhead, while the two fragmental bands after T7 endonuclease digestion are indicated by red arrowheads. Among all guides, only Ban_parn_g2 did not show successful cutting, probably due to single nucleotide polymorphisms observed in the guide region.

**Fig. S3.**
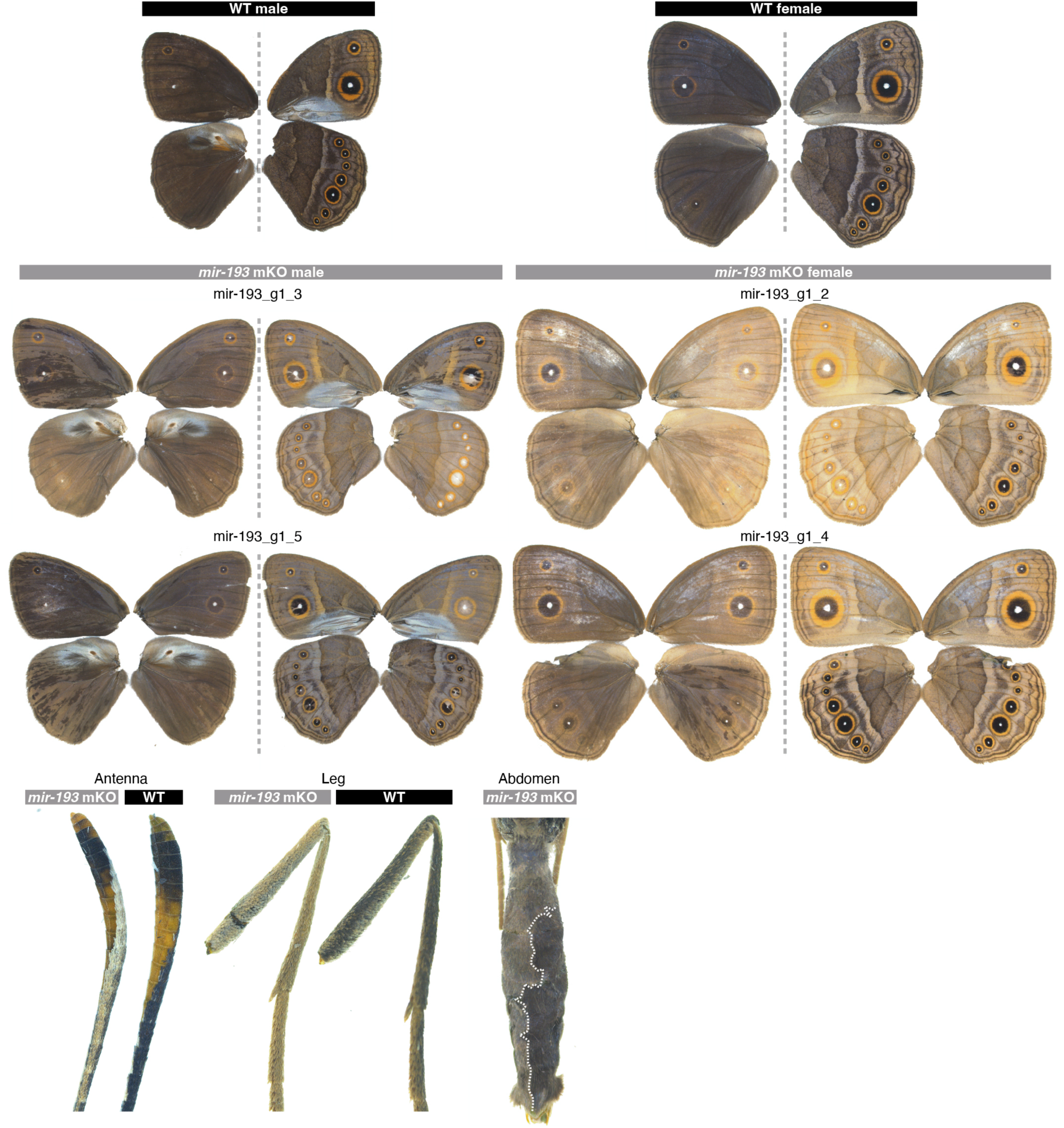
Representative phenotypes of *mir-193* mKO mutants in *B. anynana*. For the wing phenotypes, a dotted line separate dorsal (left) and ventral (right) sides of the same individual. For the abdomen, a dotted line separates the KO region and the WT region.

**Fig. S4.**
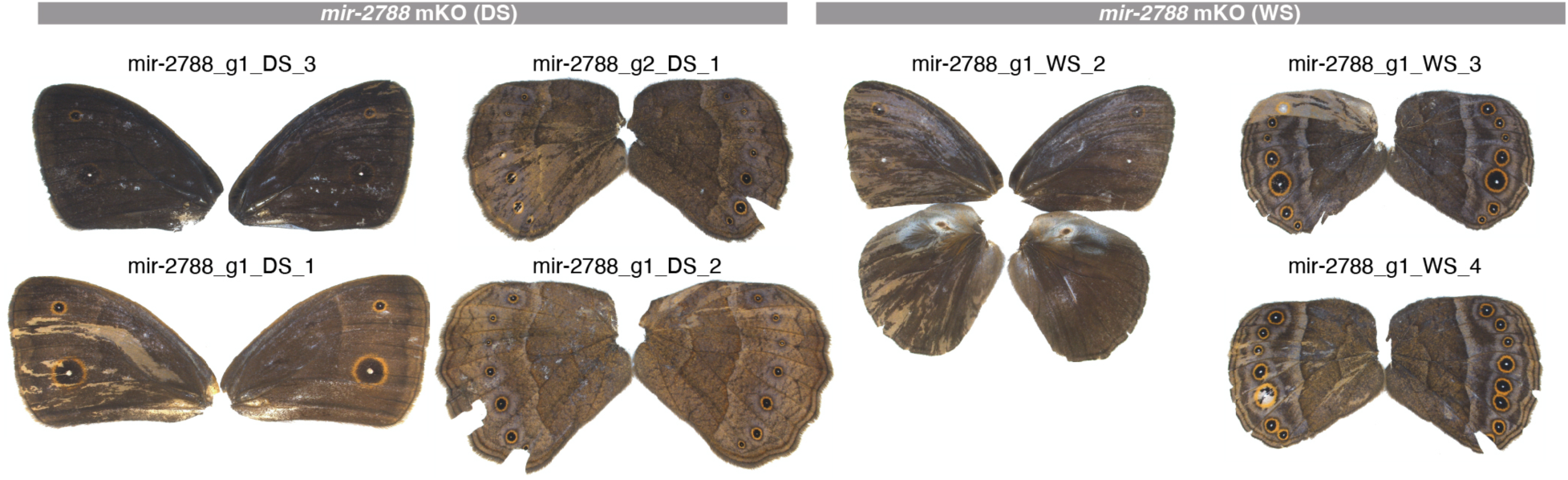
Representative phenotypes of *mir-2788* mKO mutants in *B. anynana*. In the case of *mir-2788*, injected eggs were reared either at 27°C (wet season (WS) condition), or at 17°C (dry season (DS) condition), leading to the development of the WS form with large ventral ‘eyespot’ patterns, and the DS form with small ventral ‘eyespot’ patterns, respectively.

**Fig. S5.**
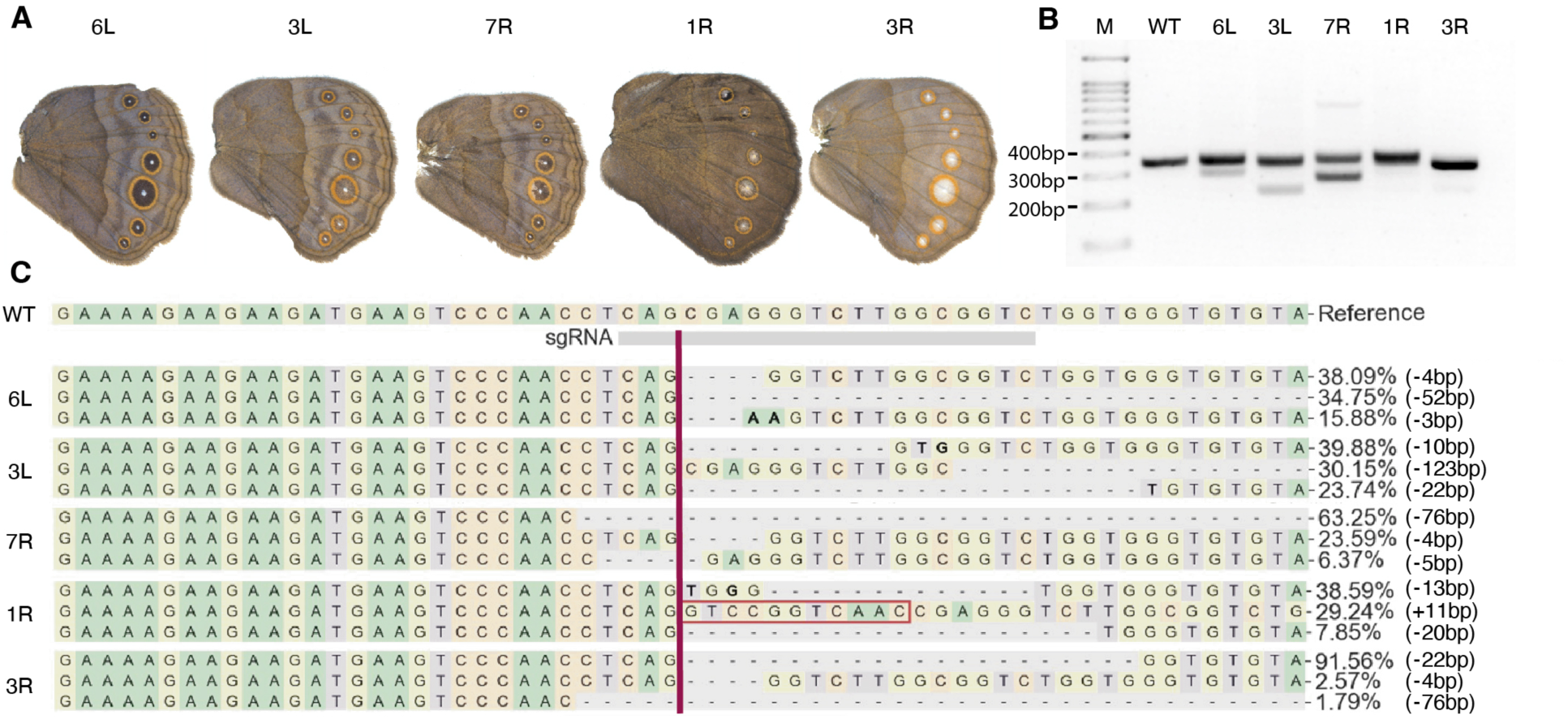
Genotyping *mir-193* mKO mutants using Illumina sequencing. **(A)** Five hindwings exhibiting a variety of melanin levels, especially in the ‘eyespot’ color pattern, from four *mir-193* mKO males were genotyped using Illumina short-read sequencing. Wings with the same number are from the same individual. L: Left wing; R: Right wing. **(B)** A gel image depicting a PCR amplicon around the *mir-193* target region, showing a variety of mutant alleles of different sizes. **(C)** Top three most abundant mutant alleles in each mutant wing detected by Illumina sequencing are shown. Figures generated from CRISPResso2. A red line denotes the cutting site. Insertions are squared in red. Wing images were horizontally flipped when necessary.

**Fig. S6.**
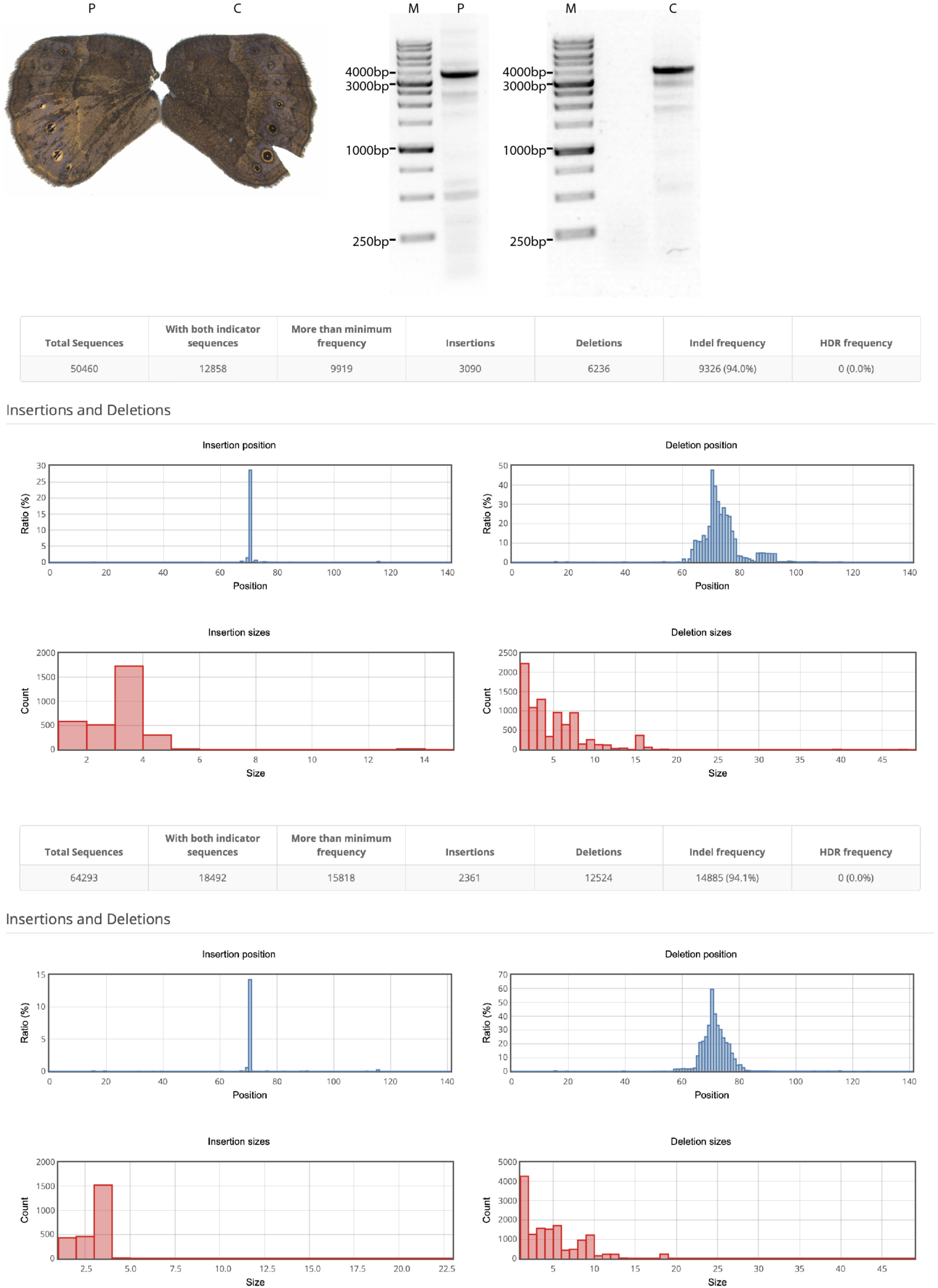
Genotyping *mir-2788* mKO mutants using Pacbio sequencing. Two hindwings from one *mir-2788* mKO male, one with reduced melanin phenotypes (P), the other without phenotypic changes (C), were genotyped using Pacbio long-read sequencing. A gel image shows the target region amplified from both wings. While confirmed long deletions were not detected in both wings (materials and methods), short on-target indels are present at a near 100% frequency in both wings, as shown in the figures generated by Cas-Analyzer.

**Fig. S7.**
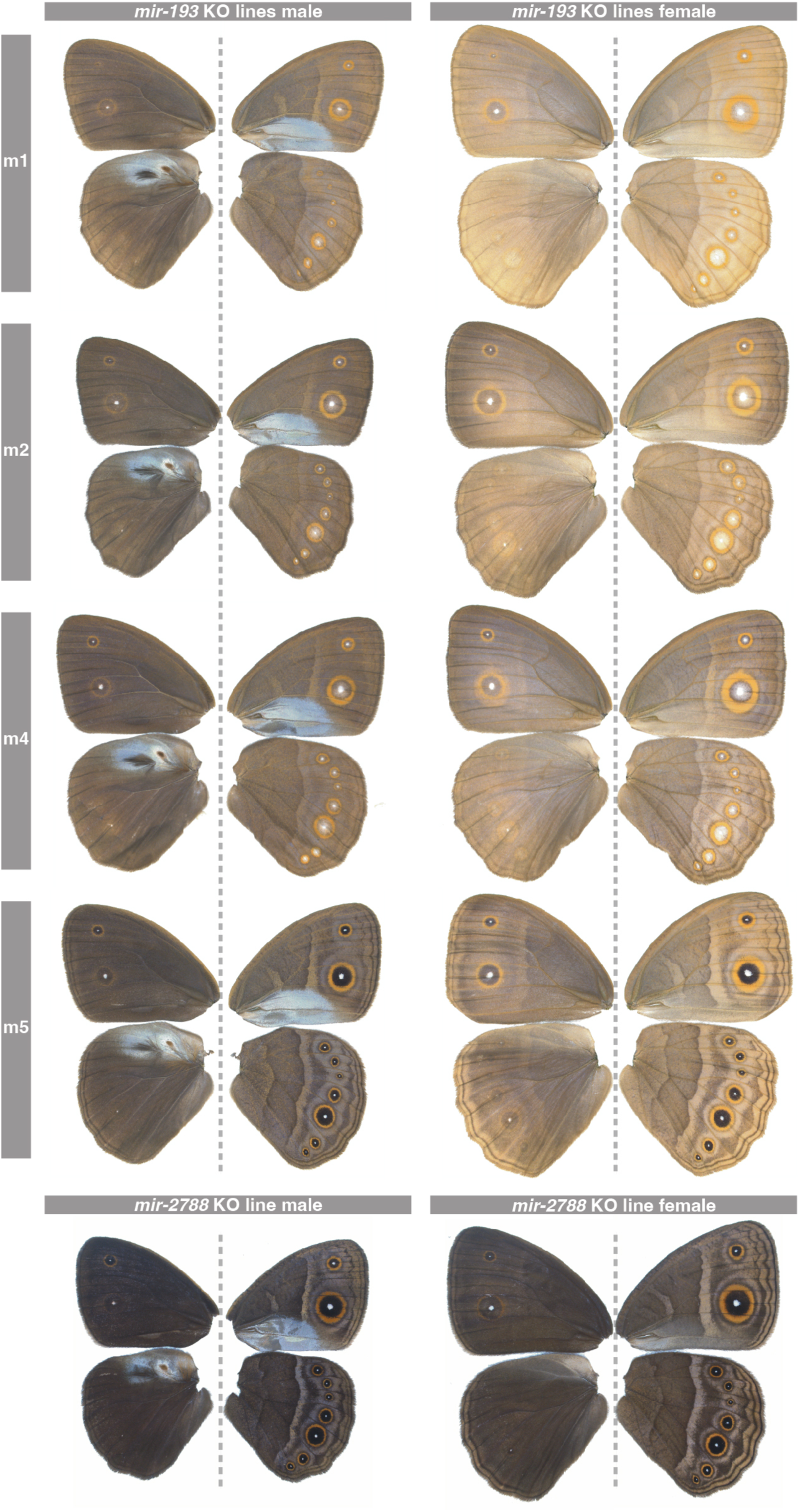
Representative phenotypes of miRNA homozygous KO lines in *B. anynana*. A dotted line separates dorsal (left) and ventral (right) sides of the same individual.

**Fig. S8.**
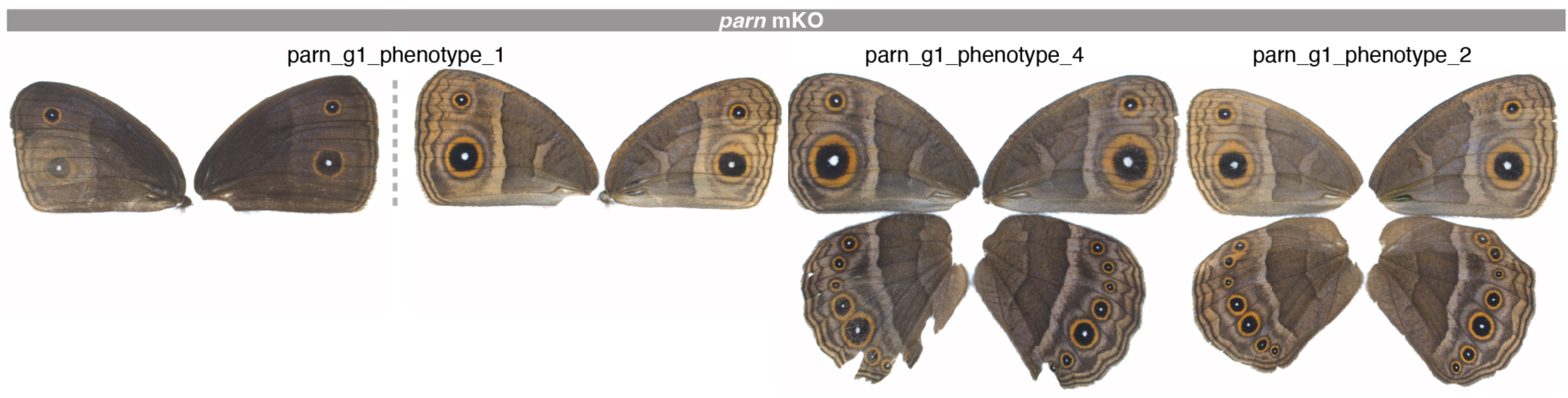
Representative phenotypes of *parn* mKO mutants in *B. anynana*. Among mKO mutants of *parn*, 19% showed reduced melanin levels on the wing that are almost invisible. A dotted line separates dorsal (left) and ventral (right) sides of the same individual.

**Fig. S9.**
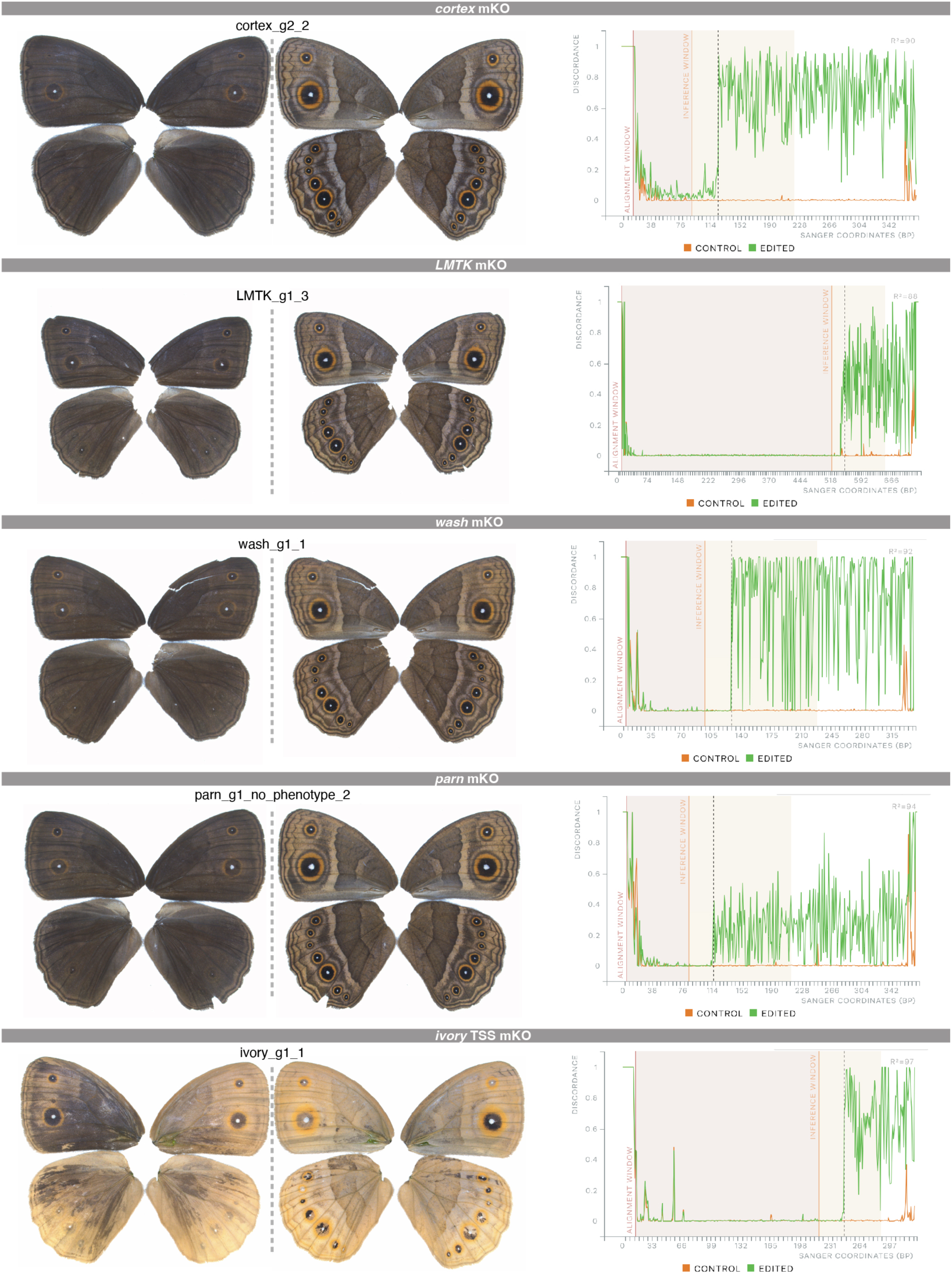
Sanger genotyping of the representative mKO mutants of the four protein-coding genes, and *ivory*, in *B. anynana*. For the wing phenotypes, A dotted line separates dorsal (left) and ventral (right) sides of the same individual. For Sanger genotyping, on-target mutations were indicated by the immediate sequence discordance after the expected gRNA cutting sites (vertical dotted lines) in the edited amplicons (green), not in the WT amplicons (orange). Figures were generated from the ICE Synthego tool.

**Fig. S10.**
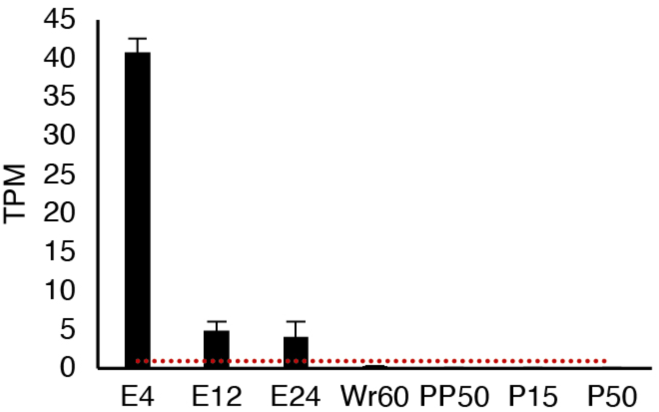
Expression levels of *cortex* across embryogenesis and wing development. Transcript per million (TPM) values of cortex across embryos 4h (E4), 12h (E12), and 24h (E24) after oviposition, and across hindwings at 60% wanderer stage (Wr60), 50% pre-pupal stage (PP50), 15% pupal stage (P15) and 50% pupal stage (P50) in *B. anynana* are shown. Detectable gene expression is defined by TPM>1 (red dotted line). Expression data is from two previous publications (*7, 15*).

**Fig. S11.**
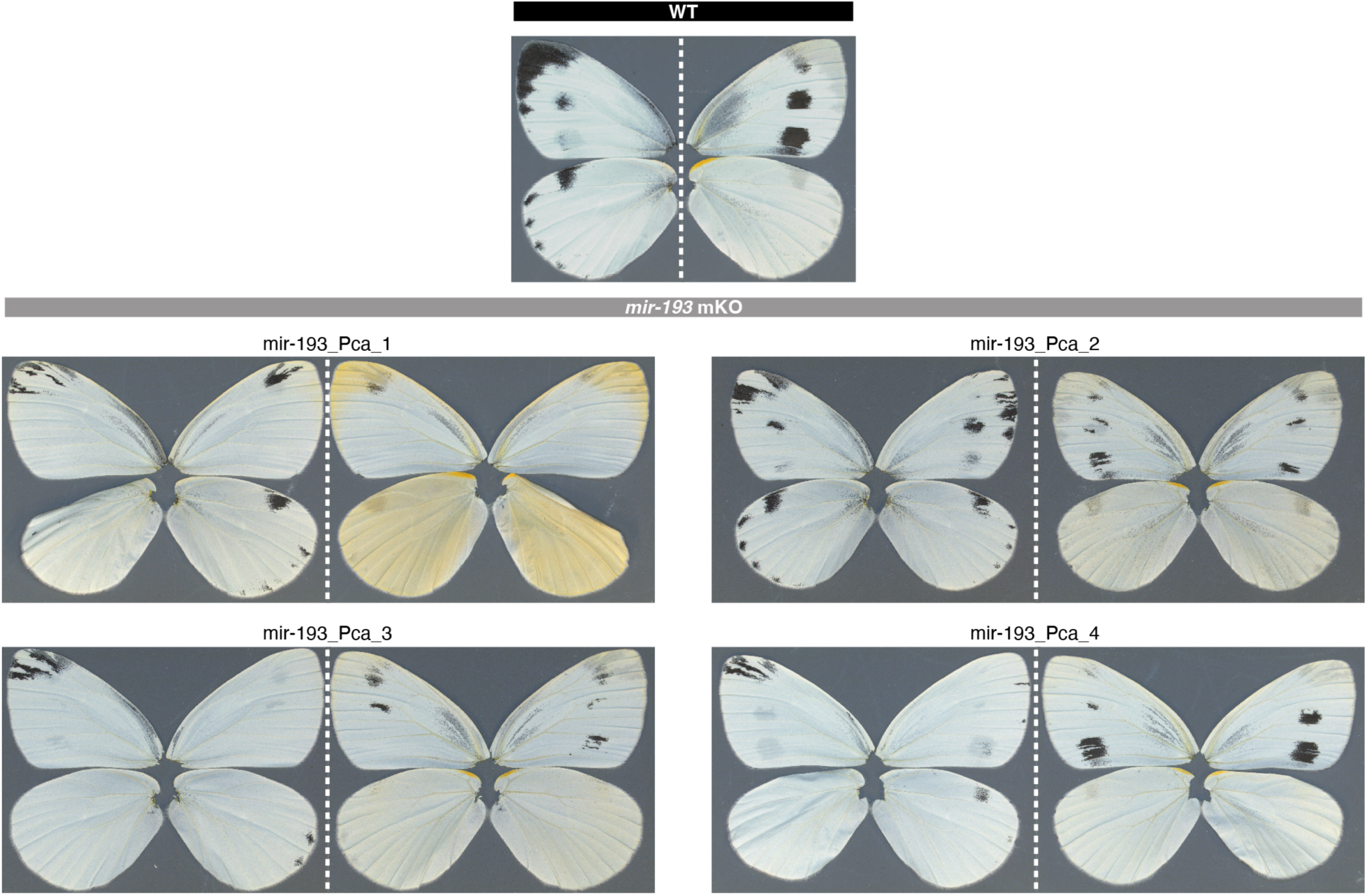
Representative phenotypes of *mir-193* mKO mutants in *P. canidia*. A dotted line separates dorsal (left) and ventral (right) sides of the same individual.

**Fig. S12.**
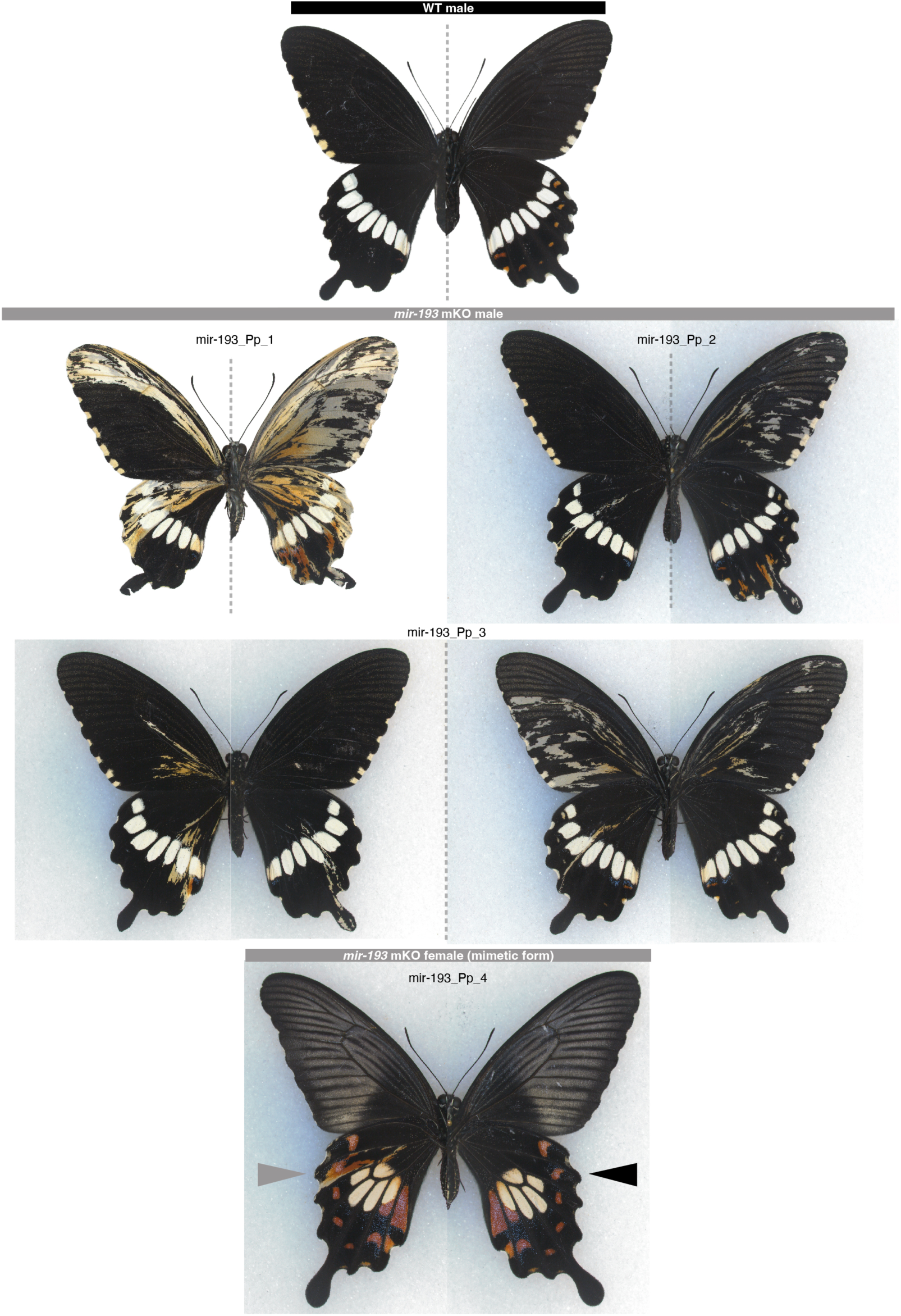
Representative phenotypes of *mir-193* mKO mutants in *P. polytes*. A dotted line separates dorsal (left) and ventral (right) sides of the same individual. In a mimetic female mutant, the edited site and unedited WT side are denoted by a grey arrowhead and a black arrowhead, respectively.

**Fig. S13.**
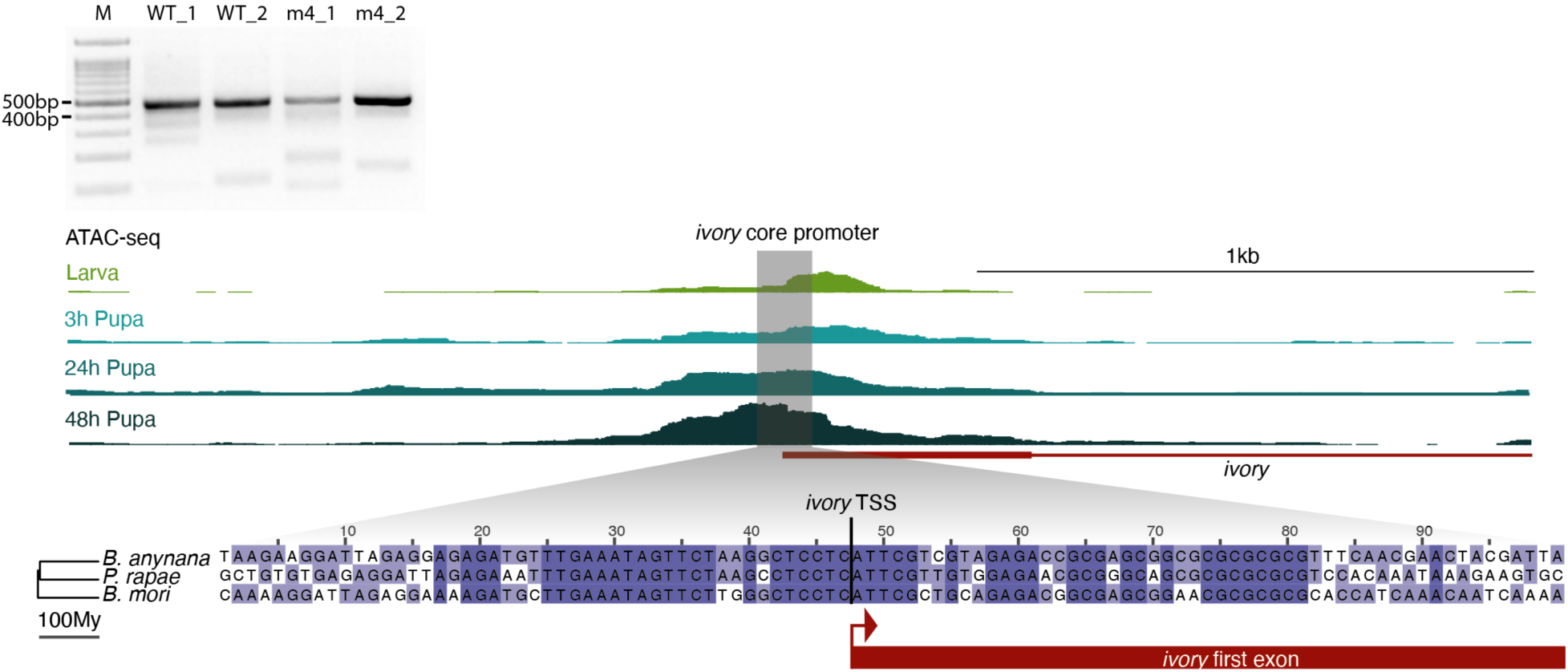
Genomic features of *ivory* TSS. To annotate *ivory* TSS, 5’RACE was performed and a consistent single band was amplified across two WT day1 pupal wings and two *mir-193* m4 mutant day1 pupal wings in *B. anynana*. Nanopore sequencing of the 5’RACE products reveals a single *ivory* TSS embedded in a 100bp promotor region deeply conserved across Lepidoptera, where an increasing chromatin accessibility can be observed during the larval-pupal transition in *B. anynana*.

**Fig. S14.**
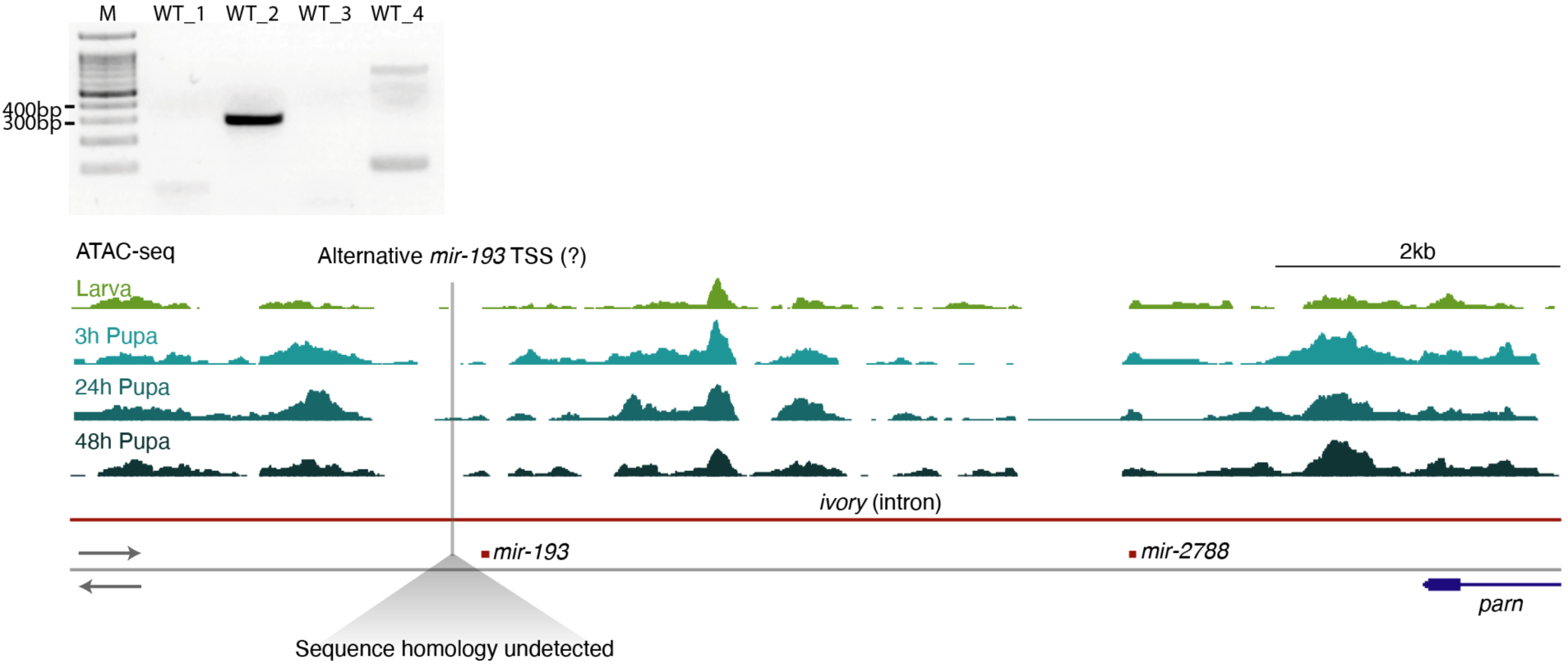
Insufficient evidence supports an alternative *mir-193* TSS independent of *ivory*. To test if an alternative TSS for *mir-193* exists independently of the *ivory* TSS, 5’RACE was performed to directly extend the *mir-193* precursor beyond its 5’ terminus from four WT day1 pupal wings of *B. anynana*. Clear 5’RACE products could only be detected from one of the four wings. Nanopore sequencing of this product reveals a putative TSS ∼200bp 5’ of *mir-193*. However, this potential TSS does not correlate with any open chromatin across wing development. Moreover, its sequence is not conserved in any other lepidopterans.

**Fig. S15.**
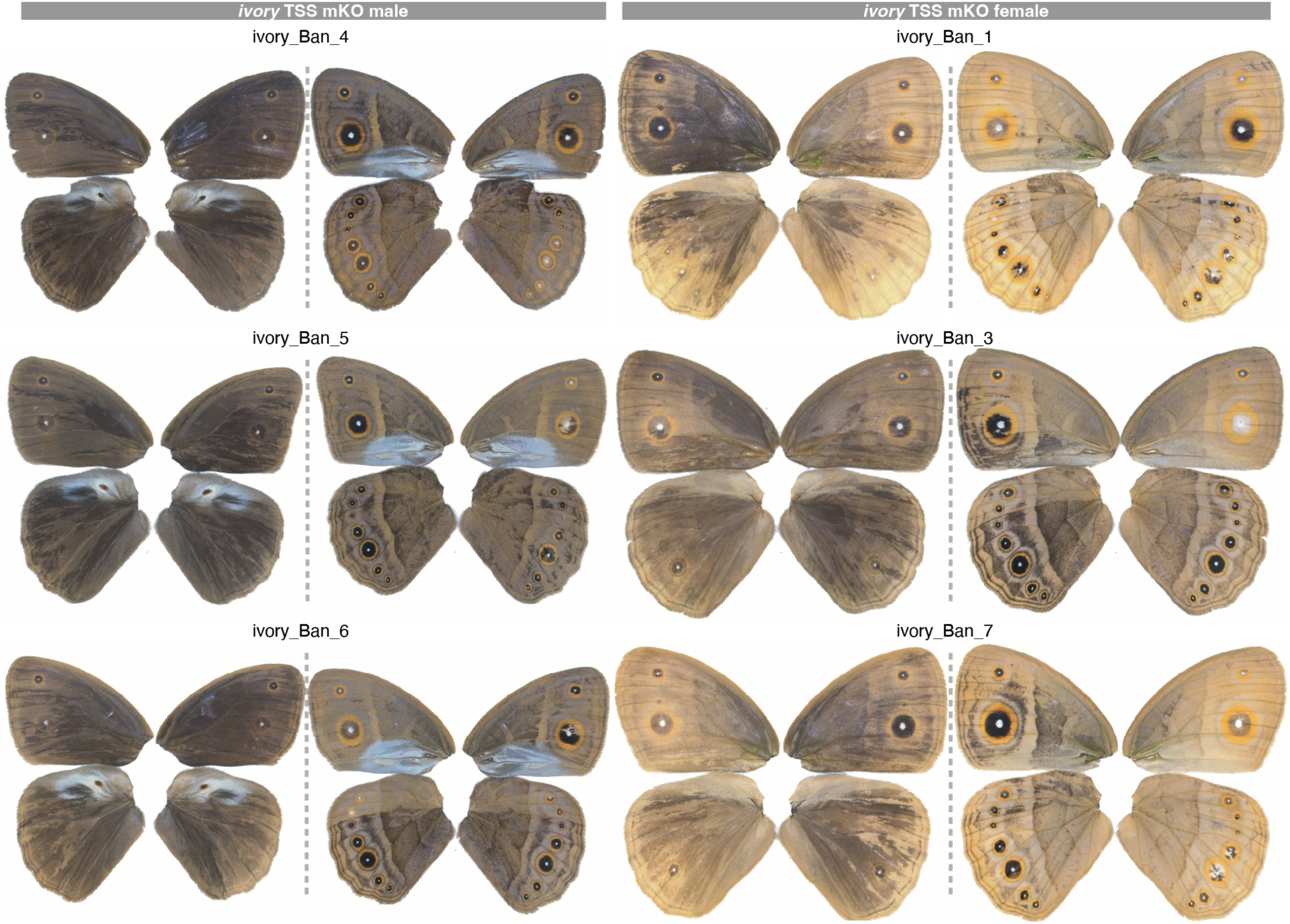
Representative phenotypes of *ivory* TSS mKO mutants in *B. anynana*. A dotted line separates dorsal (left) and ventral (right) sides of the same individual.

**Fig. S16.**
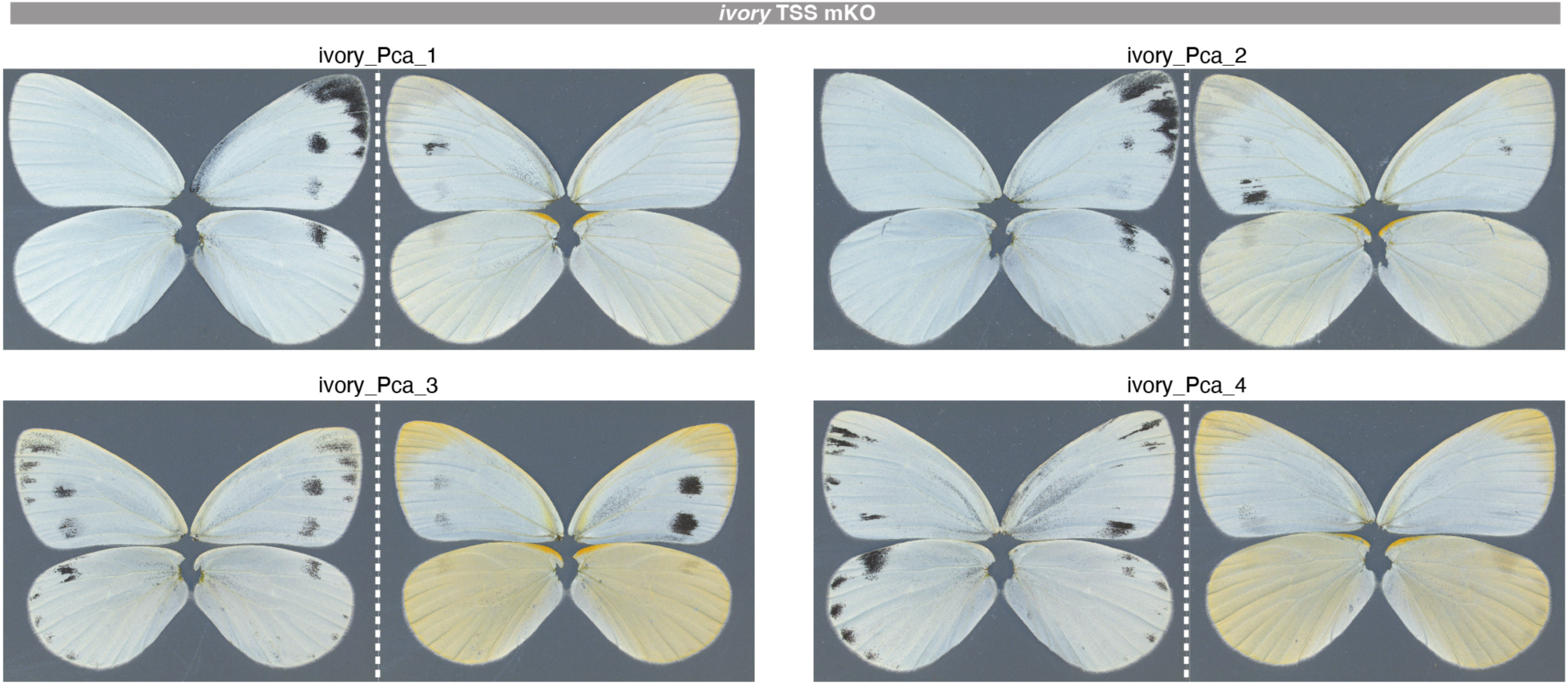
Representative phenotypes of *ivory* TSS mKO mutants in *P. canidia*. A dotted line separates dorsal (left) and ventral (right) sides of the same individual.

**Fig. S17.**
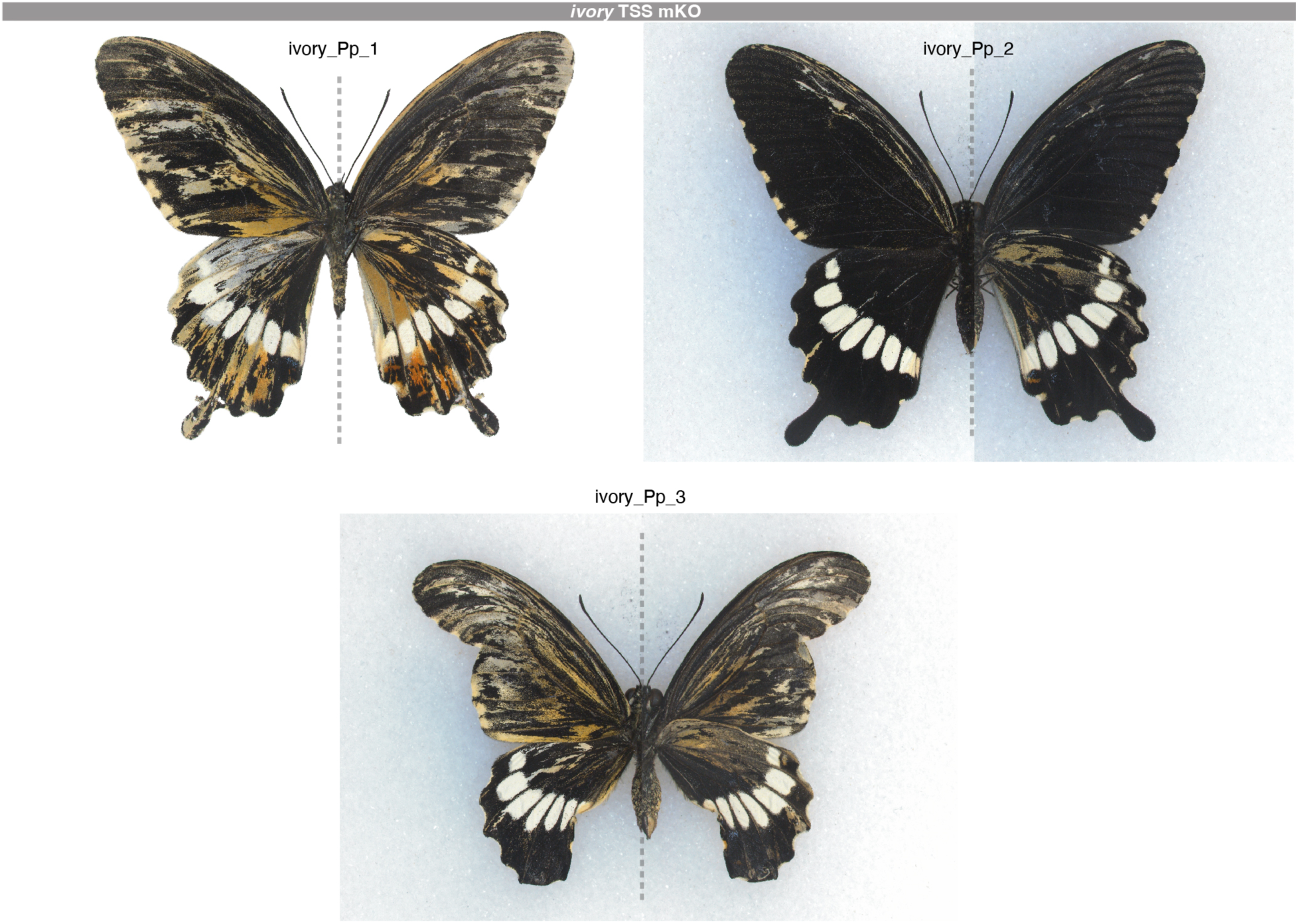
Representative phenotypes of *ivory* TSS mKO mutants in *P. polytes*. A dotted line separates dorsal (left) and ventral (right) sides of the same individual.

**Table S1.**
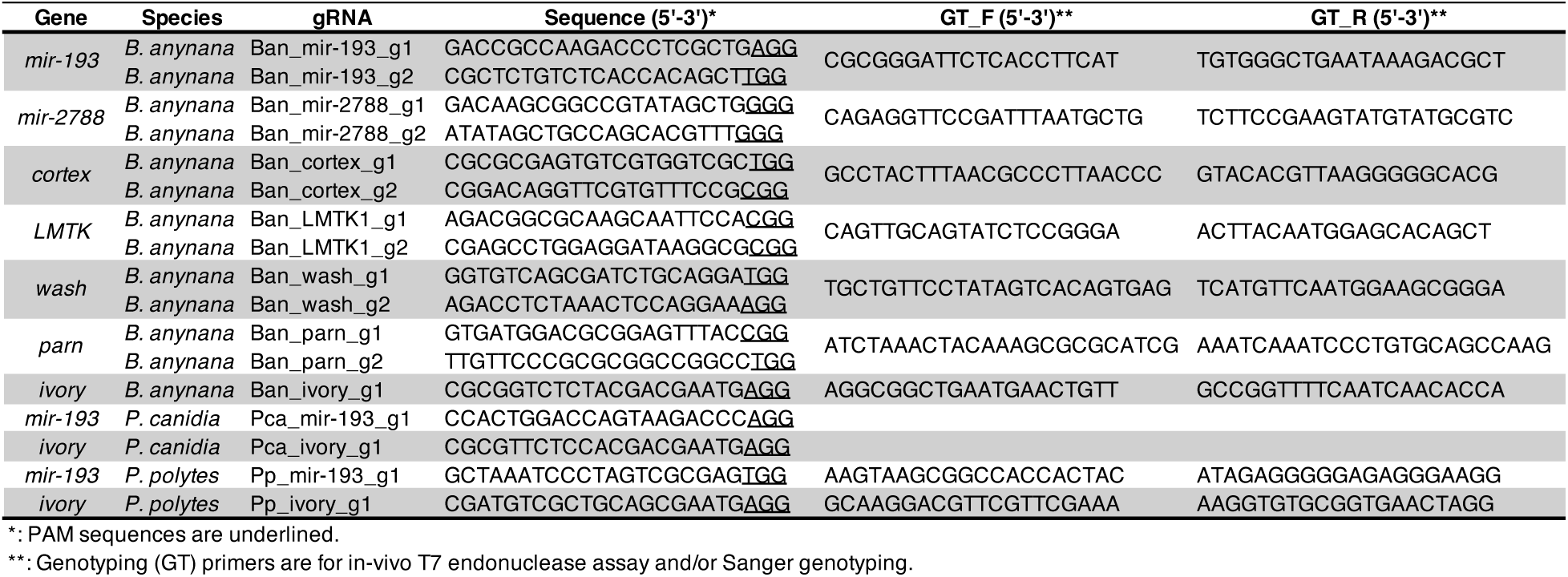
Guide RNA target sequences and genotyping primers.

**Table S2.** CRISPR injection statistics.

| Gene | Species | gRNA | Eggs injected | Rearing condition* | Hatched | % | Adults | % | Phenotypes | % |
| --- | --- | --- | --- | --- | --- | --- | --- | --- | --- | --- |
| *mir-193* | *B. anynana* | Ban_mir-193_g1 | 273 | WS | 32 | 12% | 19 | 59% | 14 | 74% |
| *mir-193* | *B. anynana* | Ban_mir-193_g2 | 277 | WS | 56 | 20% | 18 | 32% | 15 | 83% |
| *mir-2788* | *B. anynana* | Ban_mir-2788_g1 | 171 | WS | 36 | 56% | 17 | 47% | 0 | 0% |
|  |  |  |  | DS | 60 |  | 39 | 65% | 3 | 8% |
| *mir-2788* | *B. anynana* | Ban_mir-2788_g2 | 175 | WS | 20 | 33% | 12 | 60% | 0 | 0% |
|  |  |  |  | DS | 37 |  | 23 | 62% | 1 | 4% |
| *mir-2788* | *B. anynana* | Ban_mir-2788_g1 | 273 | WS | 150 | 55% | 82 | 55% | 0 | 0% |
| *mir-2788* | *B. anynana* | Ban_mir-2788_g2 | 70 | WS | 13 | 19% | 5 | 38% | 0 | 0% |
| *mir-2788* | *B. anynana* | Ban_mir-2788_g1 | 399 | WS | 140 | 35% | 90 | 64% | 6 | 7% |
| *mir-2788* | *B. anynana* | Ban_mir-2788_g1 | 356 | WS | 172 | 48% | 55 | 32% | 0 | 0% |
| *mir-2788* | *B. anynana* | Ban_mir-2788_g2 | 364 | WS | 148 | 41% | 34 | 23% | 0 | 0% |
| *cortex* | *B. anynana* | Ban_cortex_g1 | 225 | WS | 74 | 33% | 32 | 43% | 0 | 0% |
| *cortex* | *B. anynana* | Ban_cortex_g2 | 127 | WS | 32 | 25% | 24 | 75% | 0 | 0% |
| *LMTK* | *B. anynana* | Ban_LMTK1_g1 | 377 | WS | 180 | 48% | 62 | 34% | 0 | 0% |
| *LMTK* | *B. anynana* | Ban_LMTK1_g2 | 286 | WS | 118 | 41% | 65 | 55% | 0 | 0% |
| *wash* | *B. anynana* | Ban_wash_g1 | 280 | WS | 135 | 48% | 80 | 59% | 0 | 0% |
| *wash* | *B. anynana* | Ban_wash_g2 | 313 | WS | 180 | 58% | 86 | 48% | 0 | 0% |
| *parn* | *B. anynana* | Ban_parn_g1 | 375 | WS | 235 | 63% | 52 | 22% | 10 | 19% |
| *ivory* | *B. anynana* | Ban_ivory_g1 | 278 | WS | 170 | 61% | 106 | 62% | 104 | 98% |
| *mir-193* | *P. canidia* | Pca_mir-193_g1 | 137 | WS | 36 | 26% | 20 | 56% | 11 | 55% |
| *ivory* | *P. canidia* | Pca_ivory_g1 | 89 | WS | 45 | 51% | 26 | 58% | 14 | 54% |
| *mir-193* | *P. polytes* | Pp_mir-193_g1 | 197 | WS | 80 | 41% | 12 | 15% | 7 | 58% |
| *ivory* | *P. polytes* | Pp_ivory_g1 | 281 | WS | 127 | 45% | 7 | 6% | 5 | 71% |

\*: WS: wet-season condition, 27°C; DS: dry-season condition, 17°C

**Table S3.**
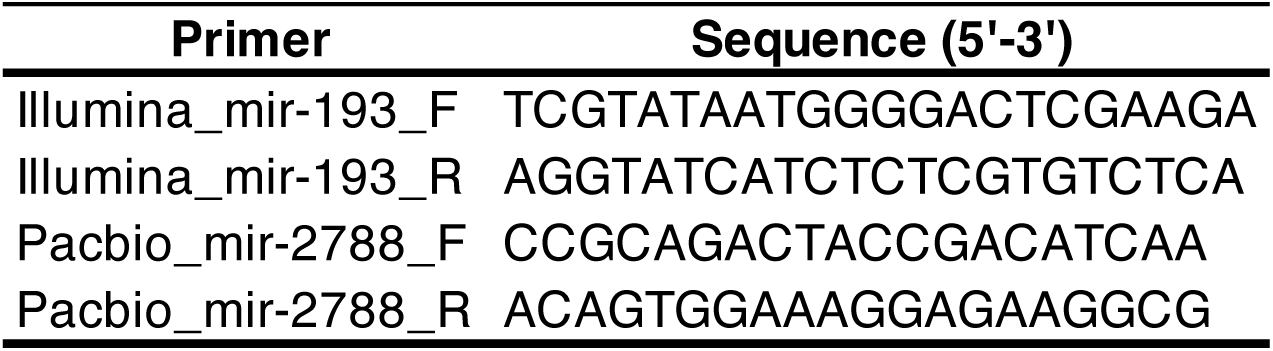
NGS genotyping primers.

**Table S4.**
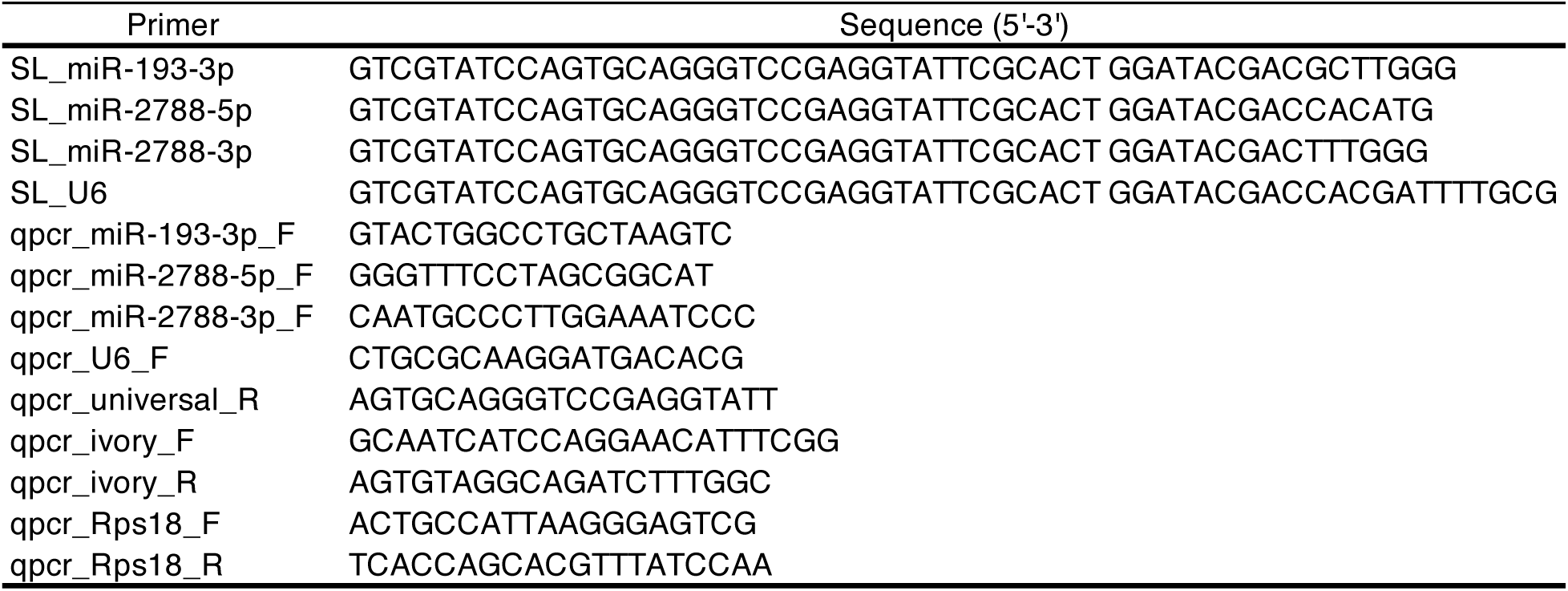
Stem-loop (SL) primers and qPCR primers.

**Table S5.** Sanger genotyping of mKO mutants in *B. anynana*.

| **Individual ID** | **Gene** | **gRNA** | **Indel rate\*** |
| --- | --- | --- | --- |
| cortex_g2_2 | *cortex* | Ban_cortex_g2 | 88% |
| cortex_g2_3 | *cortex* | Ban_cortex_g2 | 90% |
| cortex_g2_4 | *cortex* | Ban_cortex_g2 | 92% |
| cortex_g2_5 | *cortex* | Ban_cortex_g2 | 93% |
| parn_g1_1 | *parn* | Ban_parn_g1 | 17% |
| parn_g1_2 | *parn* | Ban_parn_g1 | 24% |
| LMTK_g1_3 | *LMTK* | Ban_LMTK_g1 | 60% |
| LMTK_g2_1 | *LMTK* | Ban_LMTK_g2 | 17% |
| LMTK_g2_3 | *LMTK* | Ban_LMTK_g2 | 26% |
| wash_g1_1 | *wash* | Ban_wash_g1 | 90% |
| wash_g1_3 | *wash* | Ban_wash_g1 | 63% |
| wash_g2_1 | *wash* | Ban_wash_g2 | 95% |
| wash_g2_2 | *wash* | Ban_wash_g2 | 90% |
| wash_g2_3 | *wash* | Ban_wash_g2 | 74% |
| ivory_g1_1 | *ivory* | Ban_ivory_g1 | 91% |
| ivory_g1_2 | *ivory* | Ban_ivory_g1 | 89% |

\*: Calculated from the ICE Synthego tool

**Table S6.**
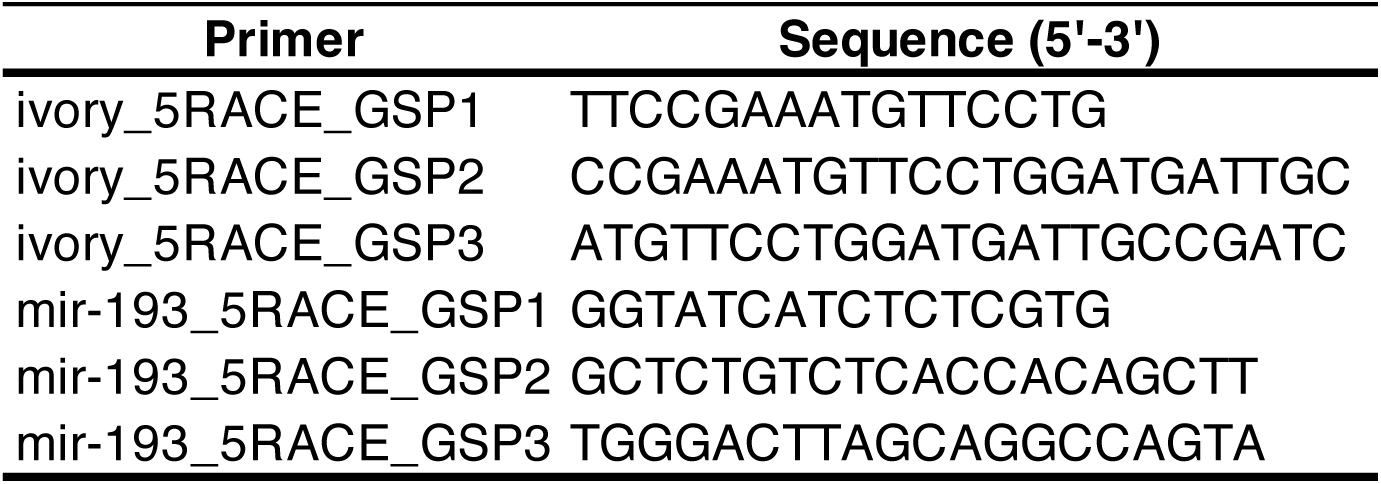
5’RACE gene-specific primers (GSPs)

**Table S7.**
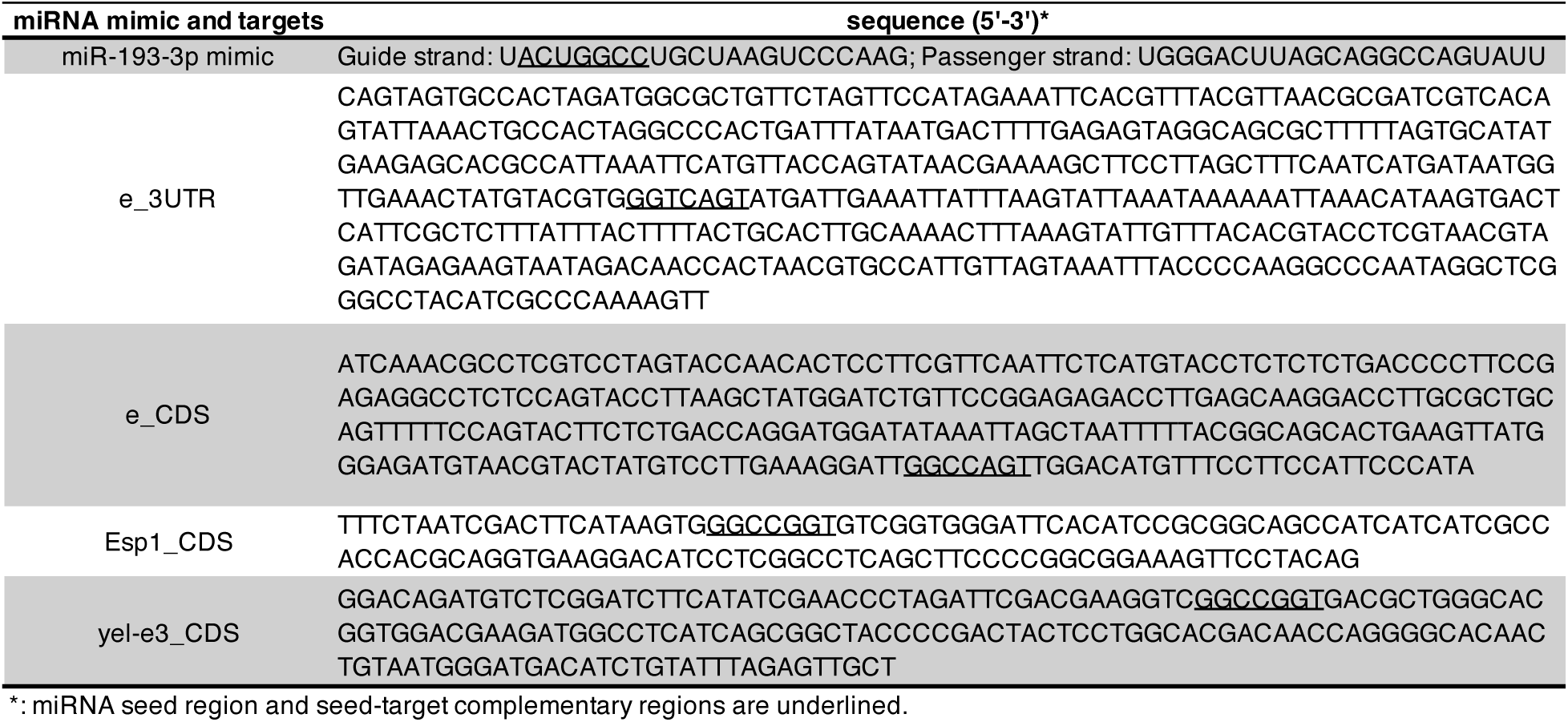
miR-193-3p mimic and dual luciferase assay target sequences.

**Data S1. (separate file)**

HCR probes

**Data S2. (separate file)**

Differentially expressed genes between *mir-193* m4 mutant and WT

**Data S3. (separate file)**

Predicted direct targets of *mir-193*

